# How does stochasticity in learning impact the accumulation of knowledge and the evolution of learning?

**DOI:** 10.1101/2025.08.26.672373

**Authors:** Ludovic Maisonneuve, Laurent Lehmann

## Abstract

Learning is crucial for humans and other animals to acquire knowledge, enhancing survival and reproduction. In particular, individual and social learning allow populations to accumulate knowledge across generations. Here, we examine how stochasticity in the production and social acquisition of knowledge influences the evolution of learning strategies and cumulative knowledge. Using a mathematical model where learning is stochastic, we show that learning stochasticity enhances cumulative knowledge by generating variability in knowledge levels. This allows selection to enhance population knowledge: individuals who acquire more knowledge by chance are more likely to survive and reproduce, and therefore to transmit their knowledge to the next generation. As knowledge accumulates, social learning exemplars tend to possess more of it, favoring greater time investment in social learning. Because social learning provides access to substantially more knowledge when learning is stochastic, selection also favors the evolution of greater investment into learning, at the expense of a fecundity cost. Moreover, when knowledge enhances fecundity but not survival, learning stochasticity favors learning from parents rather than other adults, because learning stochasticity increases uncertainty about exemplar knowledge, making parenthood a cue for possessing fecundity-enhancing knowledge. Finally, when learning occurs predominantly from parents, learning stochasticity itself is favored by selection.

## 1 Introduction

Individuals use knowledge to perform behaviors that enhance survival and reproduction. This knowledge can be acquired either through personal experience (individual learning), such as trial-and-error learning (e.g., Dugatkin, 2008; Ghirlanda and Lind, 2017), or from other individuals (social learning), for example, through imitation (e.g., Dugatkin, 2008; Zentall, 2006; Bates and Byrne, 2010). Social learning can occur through a wide variety of pathways (CavalliSforza and Feldman, 1981; Boyd and Richerson, 1985; Laland, 2004; van Schaik, 2016; Kendal et al., 2018; Camacho-Alpízar and Guillette, 2023), including, for instance, vertical transmission (from parent to offspring) and oblique transmission (from unrelated older individuals). The way individuals allocate resources to different learning behaviors across their lifetime affects the population dynamics of knowledge and can support its gradual accumulation and refinement across generations. This process, known as cumulative knowledge or cumulative culture, is widely regarded as a key factor in the ecological success of humans (Henrich, 2015; van Schaik, 2016), and growing evidence suggests that it may also contribute to increasingly adaptive behaviors in non-human animals (Hunt and Gray, 2003; Sasaki and Biro, 2017; Jesmer et al., 2018; Gunasekaram et al., 2024).

Learners’ individual and social learning strategies influence changes in knowledge within a population by shaping *cultural deviation* and *cultural selection*, the two key mechanisms underlying population change in any cultural trait and understood here as a socially transmissible trait (see Henrich and Boyd, 2002; El Mouden et al., 2014; Aguilar and Akçay, 2018; Nettle, 2020; Mesoudi, 2021). First, cultural deviation occurs when the learning process, on average, leads to differences between the knowledge of learners and that of their exemplars. This mechanism can either promote knowledge accumulation, for example, when learners produce new or refine existing knowledge by individual learning after social learning (e.g., Enquist et al., 2008; Aoki et al., 2012a; Nakahashi, 2013; Wakano and Miura, 2014; Kempe et al., 2014; André and Baumard, 2020; Denton et al., 2023), or constrain it, as when learners tend to acquire a lower level of knowledge than their exemplars (e.g., Henrich and Boyd, 2002; Henrich, 2004). Second, cultural selection arises when some individuals serve as exemplars more frequently, amplifying the transmission of their specific knowledge or cultural trait and generating variation in the transmission success of different knowledge or cultural variants (Cavalli-Sforza and Feldman, 1981; Boyd and Richerson, 1985; Micheletti, 2020). Cultural selection may arise from either (i) non-random exemplar choice or (ii) differences in survival and reproduction among individuals with different knowledge or cultural traits (though some authors use the term cultural selection specifically for mechanism (i); Cavalli-Sforza and Feldman, 1981; Mesoudi, 2011). Cultural selection is generally expected to support cumulative knowledge, as it tends to favor cultural variants that are more well-adapted to environmental conditions, for example, when individuals with more adaptive cultural variants tend to be chosen as learning models (e.g., Henrich, 2004; Powell et al., 2009; Kobayashi and Aoki, 2012), or when such variants enhance survival and fecundity, allowing them to spread by increasing opportunities for oblique (e.g., Nakahashi, 2010) and vertical transmission (e.g., Tureček et al., 2019), respectively. Mathematical models have shown that the strength of cultural selection, and thus its potential to drive cumulative knowledge, increases with population variance in cultural traits (Cavalli-Sforza and Feldman, 1981; Boyd and Richerson, 1985).

In turn, knowledge dynamics shape what individuals can acquire through social learning, thereby influencing selection pressures on the allocation of resources across different types of learning behavior. This feedback between knowledge accumulation and learning strategies gives rise to complex coevolutionary dynamics, also shaped by trade-offs between learning and other functions essential for survival and reproduction (for empirical evidence on such trade-offs, see Mery and Kawecki, 2004; Burger et al., 2008; Snell-Rood et al., 2011; Kotrschal et al., 2013; Jaumann et al., 2013; Christiansen et al., 2016; Evans et al., 2017; Padamsey and Rochefort, 2023). A large body of theoretical work has examined these dynamics and how ecological factors influence the allocation of resources to different forms of learning and the emergence of cumulative knowledge (Nakahashi, 2010; Aoki et al., 2012b; Lehmann et al., 2013; Nakahashi, 2013; Wakano and Miura, 2014; Kobayashi et al., 2015, 2016; Mullon and Lehmann, 2017; Ohtsuki et al., 2017; Maisonneuve et al., 2025). Nevertheless, the effect of cultural selection is generally neglected in these models (with few exceptions such as Kobayashi et al., 2016), often because these models assume a deterministic knowledge acquisition process at the individual level, which removes variation that can be selected upon.

However, cultural selection is likely to influence the coevolution of cumulative knowledge and learning strategies, as learning is inherently stochastic; for example, the success of trial-and-error learning often depends on chance discoveries of adaptive cultural variants, leading to individual variation in knowledge. While not their primary focus, Kobayashi et al. (2016) showed that stochasticity in individual learning promotes investment in social learning by amplifying the effect of cultural selection (caused by knowledge-based choice of social exemplars), thereby increasing overall population knowledge and increasing what can be acquired socially. However, in contrast to the assumptions of Kobayashi et al. (2016), individuals in natural populations may not always be able to reliably assess the knowledge of others and to use it to guide their social learning decisions (Argyle and McHenry, 1971; Lutz and Keil, 2002; Wood et al., 2012; Jiménez and Mesoudi, 2019; Hirel et al., 2025). This underscores the need to investigate how stochasticity in learning affects the coevolution of cumulative knowledge and learning strategies without knowledge-based choice of social exemplars.

Furthermore, there is a lack of predictions about how stochasticity in learning affects the evolution of the choice among transmission pathways (e.g., vertical vs. oblique), or the trade-offs between learning and other functions essential for survival and reproduction. Stochasticity in learning could affect these features because it increases uncertainty about the knowledge held by potential exemplars. Previous models have shown that, under such uncertainty, selection favors learning knowledge that affects fecundity from parents rather than from other adults (McElreath and Strimling, 2008). This suggests that stochasticity in learning may play an important role in shaping the allocation between vertical and oblique transmission.

In this study, we examine how stochasticity in learning influences knowledge accumulation by developing an evolutionary model in which learning is described as a stochastic process. The model tracks the evolution of the overall resource allocation to learning and fecundity, as well as the allocation of time across different types of learning (vertical, oblique, and individual learning). This allows us to examine how stochasticity influences the evolution of learning strategies, specifically, the pathways individuals use to acquire information through social learning and the trade-off between investment in learning and reproduction. By allowing knowledge to accumulate over generations, our framework also captures the coevolutionary feedback between learning strategies and knowledge accumulation.

## 2 Model

### 2.1 Life-cycle

We consider a large, asexual population where individuals acquire information that enhances fecundity and survival. This includes information such as the location of food sources, the edibility of different foods, or instructions on how to build and use a tool. Each individual possesses a quantity of adaptive information, referred to as knowledge, which is treated as a quantitative variable in our analysis. In each generation, the population goes through the following life-cycle events (see fig. 1a). (1) Adults produce offspring depending on their knowledge. (2) Offspring acquire knowledge socially from adults and through individual learning through a stochastic learning process. (3) Parents die. Offspring go through a density-dependent survival stage, where their knowledge affects survival. Those surviving become the adults of the next generation, and the cycle starts again. As a result of reproduction and survival, the number of individuals is not fixed and may vary across generations.

**Figure 1:**
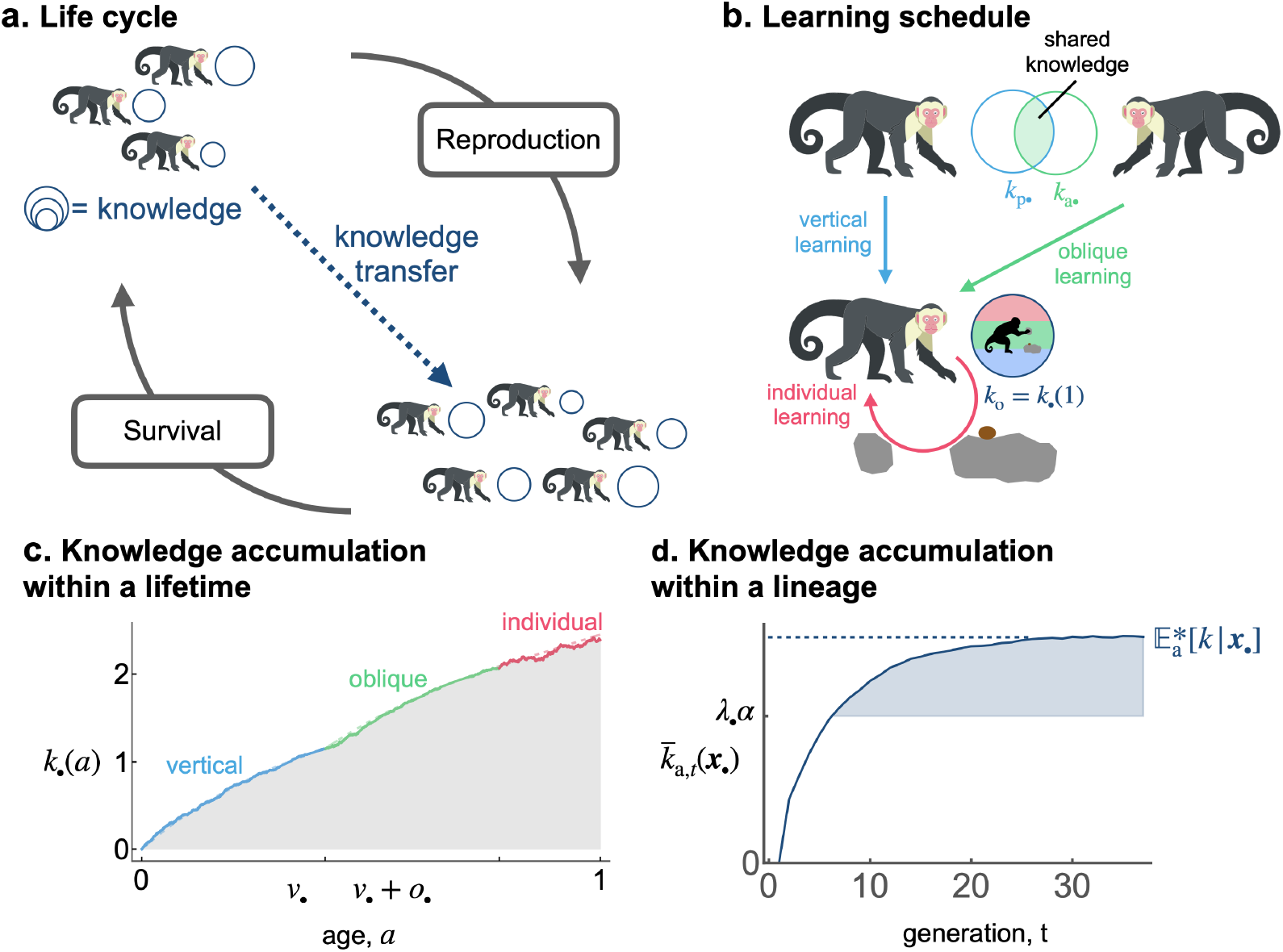
Model overview. **a** Illustration of the life cycle. **b** Illustration of the learning process. A focal individual can obtain knowledge (e.g., the skill set to crack nuts open, denoted by *k*_o•_ = *k*_•_(1) and represented here as a round set) by learning from three sources: (i) vertically from its parent (with knowledge *k*_p•_; blue arrow); (ii) obliquely from a randomly selected adult (with knowledge *k*_a•_, the knowledge of the parent *k*_p•_ and the oblique exemplar *k*_a•_ can also overlap and thus be redundant; green arrow); and (iii) individually, when it produces its own knowledge (in pink). See main text section 2.2.2 for more details. **c** A realisation of knowledge accumulation with a lifetime: individual knowledge *k*_•_(*a*) of a focal offspring against its age *a* (realisation of the stochastic process defined by eq. (2) with traits *v*_•_ = 0.4, *o*_•_ = 0.38 and *λ*_•_ = 0.82 for the offspring; and parameters *β*_v_ = 3, *β*_o_ = 2.4, *α* = 2, *ϵ* = 0.25, *ρ* = 0.05, *σ*_v_ = *σ*_o_ = 0.1, *σ*_i_ = 0.3, *k*_p•_ = *k*_a•_ = 2.45). The dashed line shows knowledge accumulation in the absence of stochasticity in learning, that is, when *σ*_v_ = *σ*_o_ = *σ*_i_ = 0. **d** Knowledge accumulation within a lineage: mean’s adult knowledge 𝔼_a,*t*_[*k* | ***x***_•_] within an ***x***_•_-lineage at each generation *t* (obtained from an individual-based simulation using the same parameters as in panel c, with trait mutation turned off and starting with a population of one ancestral individual with no knowledge, we set *k*_p•_ = *k*_a•_ = 0 for the ancestral individual, with *γ* = 0.1, *f*_0_ = 5, *s*_0_ = 1, *η*_f_ = 25, *η*_s_ = 5, *θ* = 0.5); see Appendix D for more detail on individual-based simulations). The shaded area corresponds to cumulative knowledge (where individuals, on average, possess more knowledge than they could acquire through individual learning alone, i.e., where 𝔼_a,*t*_[*k* | ***x***_•_] *> λ*_•_*α*). The dashed line shows the expected knowledge of a random adult of the ***x***_•_-lineage at equilibrium 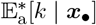 predicted by our analysis (see Section 2.3).

During stage (2) of the life cycle, offspring perform sequentially three different types of learning: first, they learn from their parent (vertical learning), then from a randomly selected adult (oblique learning), and finally by themselves (individual learning; see fig. 1b). All individuals learn during a fixed time, which we normalize to 1. Accordingly, the time allocated to vertical, oblique, and individual learning must sum to one. Two evolving traits shape how individuals allocate their time across the three different types of learning: the amount of time *v* spent learning vertically (Table 1 for a list of symbols); the amount of time *o* spent learning obliquely; so that 1 − *v* − *o* is spent learning individually.

**Table 1:**
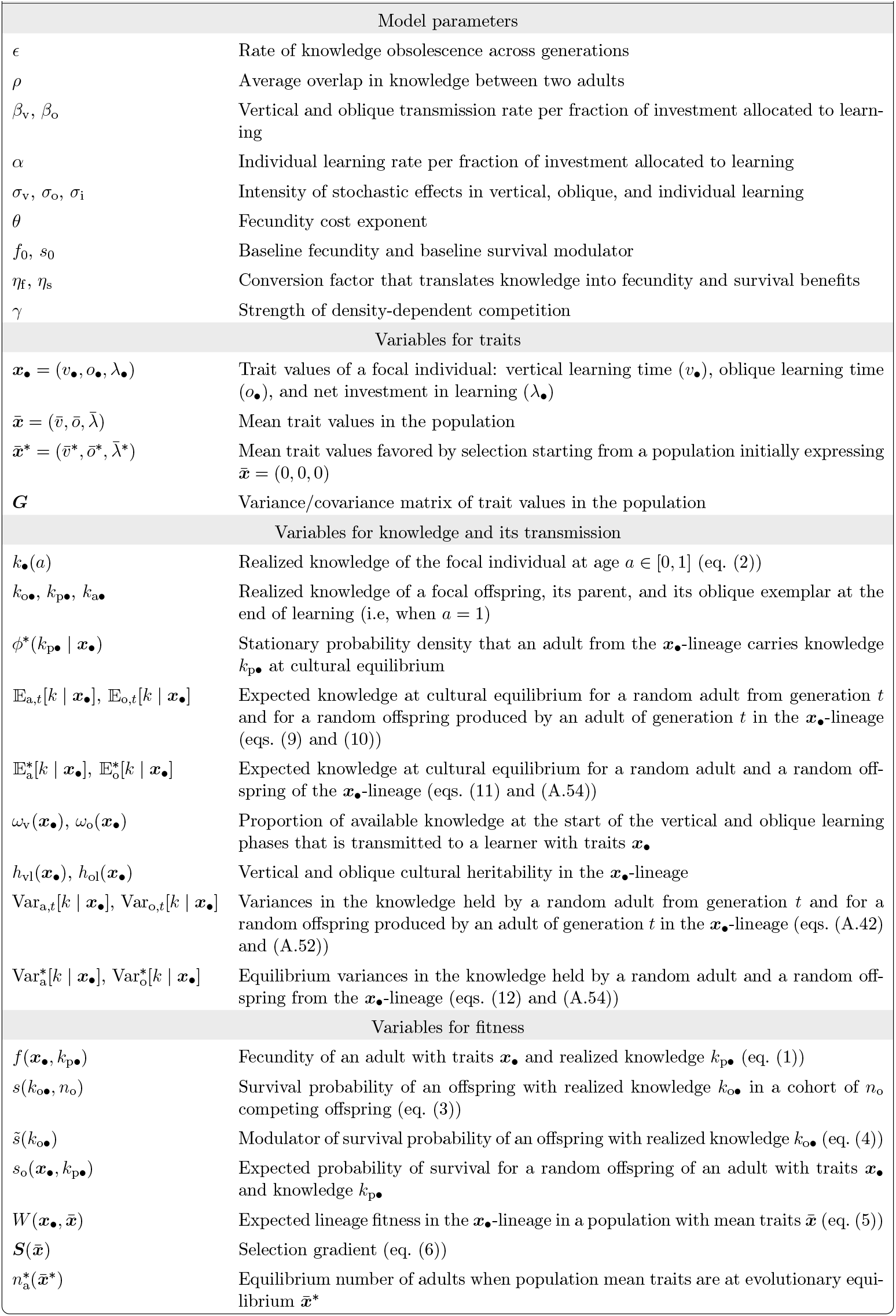
Key symbols and their definitions.

Moreover, we assume that an evolving trait *λ*∈ [0, 1] controls the overall trade-off between allocating resources to learning and fecundity. Higher values of *λ* enhance the efficiency of all learning types but at the cost of reduced fecundity. For instance, *λ* may reflect developmental costs or the metabolic expenditure associated with learning ability.

Next, we describe in detail how traits affect the three life cycle stages (Section 2.2) and then outline the approach used to analyze the cultural and the evolutionary dynamics (Section 2.3).

### 2.2 Traits effects and knowledge throughout the life-cycle

#### 2.2.1 Adult reproduction

We start by considering a focal adult from an arbitrary generation, characterized by traits ***x***_•_ =(*v*_•_, *o*_•_, *λ*_•_) and amount of knowledge *k*_p•_ ∈ R. This knowledge is the realized outcome of a stochastic learning process experienced during the individual’s offspring stage (detailed in the next section). The expected number of offspring of the focal adult, referred to as fecundity, is given by

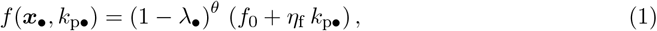

where *f*_0_ is the baseline fecundity without investment into learning, and *η*_f_ is a conversion factor translating knowledge into fecundity. For example, in a context where knowledge enhances offspring care, *η*_f_ would be high. An individual’s fecundity is reduced by its investment *λ*_•_ in learning, reflecting costs associated with acquiring learning abilities, as well as the metabolic expenses involved in the learning process itself. The parameter *θ>* 0 controls the strength and shape of this fecundity penalty: higher values of *θ* amplify the fecundity cost of learning, while lower values make this cost more gradual. This implementation of the learning–fecundity trade-off differs from earlier models, in which the trade-off between learning and reproduction is typically expressed through the allocation of time among learning and other functions (Lehmann et al., 2010, 2013; Wakano and Miura, 2014; Mullon and Lehmann, 2017).

#### 2.2.2 Offspring learning

Next, we consider an offspring of the focal adult. This focal offspring inherits the parent’s traits (*v*_•_, *o*_•_, *λ*_•_), barring mutation. Let us denote by *k*_•_(*a*) ∈ R the knowledge this offspring bears at age *a* ∈ [0, 1] (where *a* = 0 is birth). Building on the models of Kobayashi et al. (2016) and Maisonneuve et al. (2025) (where differences with our model are outlined in Appendix A.1), we model learning as a continuous-time stochastic process, in which individuals acquire knowledge at a deterministic rate on average, but this accumulation is subject to random fluctuations. Specifically, the knowledge *k*_•_(*a*) of the focal offspring at each age *a* ∈ [0, 1] is a realization of the following stochastic differential equation

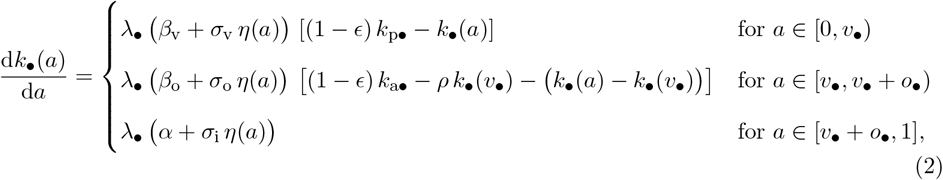

where the initial condition is *k*_•_(0) = 0. The term *η*(*a*) is the standard white noise term from stochastic calculus that introduces rapid and highly irregular fluctuations in the learning process at each age *a*, with zero mean and no temporal correlation (Gardiner, 1985, chapter 4.1). Each realization of the random white noise term *η*(*a*) over *a* ∈ [0, 1] determines a corresponding realization of knowledge acquisition *k*_•_(*a*) over *a* ∈ [0, 1] (e.g., fig. 1c).

Equation (2) says that the focal offspring first learns from its parent. Vertical learning occurs over a duration of length *v*_•_, during which the focal offspring acquires knowledge instantaneously at a rate proportional to a stochastic vertical transmission rate, *λ*_•_ (*β*_v_+*σ*_v_ *η*(*a*)), which increases with the net investment in learning *λ*_•_. The term *σ*_v_ *η*(*a*) describes the impact of stochastic events during vertical learning at age *a*, where the parameter *σ*_v_ controls the magnitude of the impact of learning stochasticity. When *σ*_v_ = 0, learning occurs with no stochastic fluctuations. The knowledge acquisition rate is assumed to be proportional to the difference (1−*ϵ*) *k*_p•_ −*k*_•_(*a*), between the knowledge currently available (1 − *ϵ*) *k*_p•_ from the parent and the focal offspring’s knowledge *k*_•_(*a*), where *ϵ* ∈ [0, 1] is the proportion of knowledge that becomes obsolete between two generations. Indeed, as the amount of available knowledge decreases, the focal offspring is less likely, on average, to encounter new information during interactions with its parent, thereby slowing the learning process.

Secondly, the focal offspring learns from a random adult, whose knowledge is denoted by *k*_a•_, for a duration of length *o*_•_. Similar to the parent’s knowledge, *k*_a•_ results from a realization of the stochastic learning process undergone by the oblique exemplar in the previous generation (see details below). During oblique learning, the instantaneous knowledge acquisition rate is proportional to the product of a stochastic oblique transmission rate, *λ*_•_ (*β*_o_ + *σ*_o_ *η*(*a*)), and of the amount of knowledge held by the oblique exemplar that the focal offspring has yet acquired, which is assumed to be given by (1 − *ϵ*) *k*_a•_ − *ρ k*_•_(*v*_•_) − (*k*_•_(*a*) − *k*_•_(*v*_•_)) . This expression accounts for both the overlap between the parent’s and the exemplar’s knowledge, and the knowledge the offspring has already acquired obliquely. This expression assumes that, on average, the knowledge held by two adults in the population overlaps by a proportion *ρ* ∈ [0, 1]. Consequently, the knowledge of the parent and of the oblique exemplar also overlaps by the same proportion *ρ*. At the end of the vertical learning phase (i.e., at age *v*_•_), the focal offspring has acquired from its parent a quantity of knowledge *k*_•_(*v*_•_), of which a quantity *ρ k*_•_(*v*_•_) is also known by the oblique exemplar. The total amount of knowledge that the focal offspring can potentially acquire from the oblique exemplar is therefore (1 − *ϵ*) *k*_a•_ − *ρ k*_•_(*v*_•_). At age *a* ≥ *v*_•_, the focal offspring has already acquired from the oblique exemplar a quantity of knowledge *k*_•_(*a*) −*k*_•_(*v*_•_), which is precisely the amount gained beyond what was acquired from the parent. Hence, the remaining knowledge available from the oblique exemplar at age *a* ≥ *v*_•_ is (1 −*ϵ*) *k*_a•_ −*ρ k*_•_(*v*_•_) − (*k*_•_(*a*) −*k*_•_(*v*)) . Note that the average overlap in knowledge between two adults, *ρ*, may reflect environmental features. For instance, environmental heterogeneity can lead individuals to produce different knowledge that addresses different ecological challenges, thereby reducing overlap.

Thirdly, the focal offspring learns individually at an instantaneous stochastic rate proportional to *λ*_•_ (*α* + *σ*_i_ *η*(*a*)) for the remaining 1 − *v*_•_ − *o*_•_ time. The parameter *σ*_i_ captures the amplitude of stochastic variation in individual learning, reflecting, for example, fluctuations in attention or intrinsic randomness in the mechanistic processes underlying learning, such as trial-and-error. During each learning phase, the rate of knowledge acquisition is proportional to the offspring’s net investment in learning *λ*_•_. As a result, when there is no such investment (i.e., *λ*_•_ = 0), the focal offspring does not acquire any knowledge at any age (i.e., ∀*a* ∈ [0, 1], *k*_•_(*a*) = 0). Due to stochasticity, the learning rate may occasionally become negative during each learning phase, implying that the offspring may lose knowledge. This could reflect situations where miscommunication or exploration leads to confusion, or replacement of previously held information by incorrect alternatives.

Note that both the knowledge of the parent *k*_p•_ and that of the oblique exemplar *k*_a•_ of the focal offspring result from realizations of the stochastic learning process experienced by those individuals in the previous generation. Specifically, their knowledge is obtained as an instantiation of *k*_•_(1), where *k*_•_(*a*) denotes a realization at age *a* of the stochastic differential equation defined in eq. (2), with the values of *k*_p•_ and *k*_a•_ on the right-hand side corresponding to the knowledge values of the parent’s own parent and oblique exemplar, and, in the case of the focal’s oblique exemplar, to those of its own learning exemplars. Since both reproduction and survival are affected by knowledge, the realized parental knowledge *k*_p•_ and oblique exemplar knowledge *k*_a•_ among the offspring are not solely determined by the stochastic learning process defined in eq. (2), but also by reproduction and survival, and thus by offspring survival, which we next specify.

#### 2.2.3 Offspring survival

After completing their learning, the offspring enter a density-dependent survival stage. Let *k*_o•_ = *k*_•_(1) denote the knowledge acquired by the focal offspring at the end of the learning phase, where *k*_•_(1) is the outcome of eq. (2). The probability of survival of the focal offspring is given by

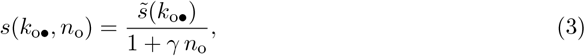

where *n*_o_ is the total number of offspring produced by all adults and *γ >* 0 is a parameter controlling the strength of density dependence. The numerator 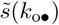, which modulates survival probability, is given by

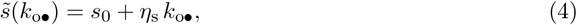

where *s*_0_ is the baseline survival, and *η*_s_ is the conversion factor that translates knowledge into survival. For example, in a context where knowledge enables predator recognition, *η*_s_ would be high. Parameter values are chosen in our analyses such that *s*(*k*_o•_, *n*_o_) remains between 0 and 1 for all individuals.

### 2.3 Analyses

Here, we detail the hypotheses and the method we employ to investigate the joint cultural and evolutionary dynamics. Assuming a small mutation rate and a large population size, evolutionary change proceeds more slowly than cultural dynamics. This timescale separation allows us to analyze cultural dynamics while treating population trait values as constant and consider the knowledge dynamics within lineages of individuals bearing the same trait values (Mullon and Lehmann, 2017).

#### 2.3.1 Cultural dynamics

We first aim to describe the equilibrium probability density of knowledge for a member of an ***x***_•_-lineage, which is shaped both by the stochastic learning process defined in eq. (2), which enables both the production and intergenerational accumulation of knowledge, and by the effects of knowledge on fecundity and survival given in eqs. (1) and (3), since individuals who survive and reproduce are more likely to transmit their knowledge. In general, the learning process is too complicated to obtain an explicit expression for the probability density of knowledge. To make the analysis tractable, we assume that the outcomes of social learning are deterministic, in contrast to individual learning, where we assume that producing knowledge is inherently stochastic (i.e., *σ*_v_ = *σ*_o_ = 0, *σ*_i_ *>* 0). This assumption is relaxed in individual-based simulations, which allow stochasticity in social learning and recover the same qualitative effects of stochasticity in learning on knowledge accumulation and traits evolution (fig. S.10). Under this assumption, we can solve the stochastic differential equation (2) (see Appendix A.2). Using this solution, along with the effects of knowledge on fecundity and survival given in eqs. (1) and (3), we derive a recursion for the expected knowledge and its variance for a member of the ***x***_•_-lineage (see Appendices A.3.1 and A.3.2). However, this recursion ultimately depends on the entire hierarchy of moments of the probability density of knowledge. To address this issue and be able to track the expected knowledge and its variance across generations, we employ a Gaussian closure approximation (see Appendix A.3.3). Note that the stationary distributions obtained from individual-based simulations suggest that a Gaussian approximation provides a good fit for the probability density of knowledge (see fig. S.1). With this, we can then fully characterize the cultural equilibrium in terms of the expected knowledge and its variance (see Appendix A.4).

#### 2.3.2 Evolutionary dynamics

Under the assumptions of small mutation rate, small variance in knowledge, and a large population, the expected evolutionary dynamics can be inferred from the expected lineage fitness in a focal ***x***_•_-lineage, i.e., the expected fitness of a random individual in that lineage (Mullon and Lehmann, 2017). We show in Appendix B.1 that the expected lineage fitness in the ***x***_•_-lineage in a population with mean traits 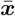 can be expressed as

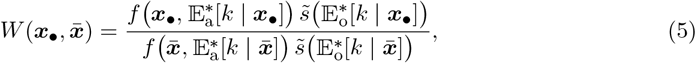

where 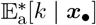 and 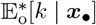 are the expected knowledge after learning is complete and at cultural equilibrium for a random adult and a random offspring of the ***x***_•_-lineage. As we assume small variance in knowledge, 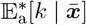 and 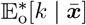 coincide with the population mean adult and offspring knowledge.

The expected evolutionary dynamics can then be inferred from the selection gradient, defined as

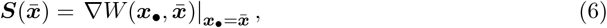

where the operator ∇ acts such that, for any function of traits *u*, we have

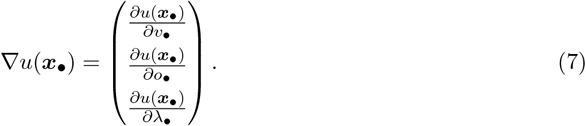

Each entry of the selection gradient indicates whether natural selection favors an increase or a decrease in the corresponding trait. According to invasion analysis (Leimar, 2009), the mean trait values 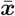 will eventually converge to a convergence stable trait vector denoted by 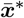, which either satisfies 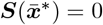 or lies on the boundary of the phenotypic space. We use the expression of the selection gradient to highlight the distinct selective pressures acting on each learning trait and to numerically estimate the trait values 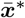 favored by selection starting from a population initially lacking any form of learning, that is, with ancestral traits 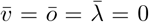 (see details in B.3). We thus focus only on the convergence stable trait vector reached from these ancestral trait values, as this corresponds to the biologically relevant scenario in which learning must first evolve before cultural dynamics operate. At 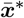, we systematically verify if selection is stabilizing (see details in B.3), ensuring the maintenance of a unimodal trait distribution. In all analyses, selection was found to be stabilizing.

To assess the robustness of our forthcoming analytical findings, we perform individual-based simulations that relax the previously stated assumptions (see details of individual-based simulations in Appendix D). In particular, we allow for small population sizes, allow *σ*_v_ *>* 0 and *σ*_o_ *>* 0, and no longer assume that the probability density of knowledge is normal.

## 3 Results

### 3.1 Cultural dynamics

#### 3.1.1 Cultural selection enhances cumulative knowledge

To reveal the effects of cultural selection on cumulative expected knowledge, we analyze the dynamics of knowledge in a given lineage. To this end, we first determine the realized knowledge *k*_o•_ acquired by a focal offspring in an ***x***_•_-lineage, whose parent has knowledge *k*_p•_ and who chooses an oblique exemplar with knowledge *k*_a•_. We show in Appendix A.2 that

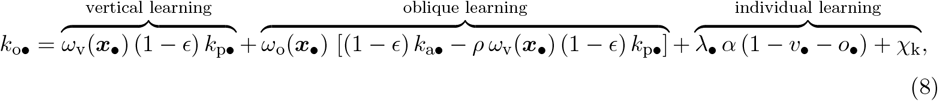

where 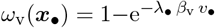 and 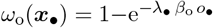 are the proportion of available knowledge at the start of the vertical and oblique learning phases, respectively, that the focal offspring acquires. Each term in eq. (8) corresponds to the amount of knowledge acquired through each type of learning. The knowledge gained through vertical learning reduces the knowledge available through oblique learning, since parental knowledge overlaps with that of the oblique exemplar with proportion *ρ*. As the offspring already acquired an amount *ω*_v_(***x***_•_) (1 − *ϵ*) *k*_p•_ of knowledge from its parent, the amount of non-redundant knowledge available at the start of oblique learning is (1 − *ϵ*) *k*_a•_ − *ρ ω*_v_(***x***_•_) (1 − *ϵ*) *k*_p•_. Finally, the knowledge acquired through individual learning *λ*_•_ *α* (1−*v*_•_ −*o*_•_)+*χ*_k_ depends on a realization of a Gaussian random variable *χ*_k_. This random variable has mean zero and variance 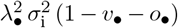, which is proportional to the time allocated to individual learning 1 − *v*_•_ − *o*_•_.

Having characterized the realized knowledge acquired by a focal offspring, we can now characterize the lineage-level dynamics, at which cultural selection acts. Specifically, by taking the expectation in eq. (8), we show in Appendix A.3.1 that the expected knowledge 𝔼_o,*t*_[*k* | ***x***_•_] of an offspring born to an adult of generation *t* in the ***x***_•_-lineage, when the expected value and variance of knowledge among adults are 𝔼_a,*t*_[*k* | ***x***_•_] and Var_a,*t*_[*k* | ***x***_•_], is given by

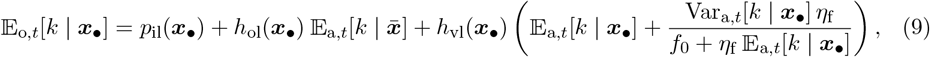

where *p*_il_(***x***_•_) = *λ*_•_ *α* (1 − *v*_•_ − *o*_•_), is the expected amount of knowledge produced through individual learning by the focal offspring, and *h*_ol_(***x***_•_) = (1 − *ϵ*) *ω*_o_(***x***_•_) and *h*_vl_(***x***_•_) = (1 − *ϵ*) *ω*_v_(***x***_•_) (1 − *ρ ω*_o_(***x***_•_)) are the oblique and vertical cultural heritabilities, respectively. The expressions for *h*_vl_(***x***_•_) and *h*_ol_(***x***_•_) are obtained by identifying, in eq. (8), the coefficients multiplying parental knowledge *k*_p•_ and oblique exemplar knowledge *k*_a•_ in the realized focal offspring knowledge *k*_o•_. The term −*ρ ω*_o_(***x***_•_) in *h*_vl_(***x***_•_) captures that, when parental and oblique knowledge overlap, knowledge acquired from the parent reduces the amount of knowledge that remains available through oblique learning. This interference effect reduces vertical, but not oblique, cultural heritability, because it scales with the amount of parental knowledge (see eq. (8)). Intuitively, this term corrects the vertical heritability *h*_vl_(***x***_•_) by accounting that a part of the knowledge acquired from the parent substitutes for knowledge that would otherwise have been acquired from oblique exemplars. The oblique and vertical cultural heritabilities, both lying between 0 and 1, can then be interpreted as the proportion of knowledge from the oblique and vertical exemplar that is effectively transmitted to a learner with traits ***x***_•_.

The last two terms on the right-hand side of eq. (9) are the expected knowledge acquired from an oblique exemplar and from the parent. These terms depend on the corresponding cultural heritabilities, multiplied by the expected knowledge 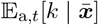 of an oblique exemplar and that 𝔼_a,*t*_[*k* | ***x***_•_] + Var_a,*t*_[*k* | ***x***_•_] *η*_f_ */* (*f*_0_ + *η*_f_ 𝔼_a,*t*_[*k* | ***x***_•_]) of a vertical exemplar. The expected knowledge of a vertical exemplar exceeds that of an average adult in the lineage, since individuals with greater knowledge produce more offspring and are overrepresented among vertical exemplars.

The term *h*_vl_(***x***_•_) Var_a,*t*_[*k* | ***x***_•_] *η*_f_ */* (*f*_0_ + *η*_f_ 𝔼_a,*t*_[*k* | ***x***_•_]) in (9) can be thought of as the response of expected knowledge to cultural selection (in analogy with the response to selection due to genetic inheritance; Lynch and Walsh, 1998), due to differences in fecundity. Because individuals with greater knowledge tend to have higher fecundity, they are more likely to transmit their knowledge vertically, which biases transmission toward higher knowledge individuals and increases the expected knowledge within the lineage. The response to cultural selection is particularly pronounced when parents transmit a substantial portion of their knowledge to their offspring (i.e., *h*_vl_(***x***_•_) is great), there is significant knowledge difference between adults (i.e., Var_a,*t*_[*k* | ***x***_•_] is great), and knowledge strongly increases fecundity (i.e., *η*_f_ is great). The strength of the response to cultural selection diminishes as relative differences in fecundity with the lineage become less pronounced with increasing values of *f*_0_ + *η*_f_ 𝔼_a,*t*_[*k* | ***x***_•_].

Following learning, the expected knowledge within a lineage is further shaped by survival to adulthood. After completing their learning, offspring go through the survival stage to reach adulthood in generation *t* + 1. We show in Appendix A.3.1 that the expected knowledge 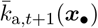 of an adult of generation *t* + 1 in the ***x***_•_-lineage is

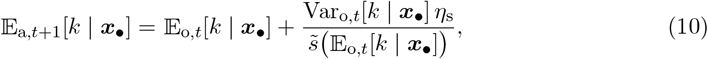

where Var_o,*t*_[*k* | ***x***_•_] is the variance in the knowledge held by a randomly chosen offspring born to an adult of generation *t* in the ***x***_•_-lineage. The expected knowledge among adults is higher than that among offspring (i.e., 𝔼_a,*t*+1_[*k* | ***x***_•_] *>* 𝔼_o,*t*_[*k* | ***x***_•_]) since offspring with above-average knowledge are more likely to survive to adulthood. This effect is captured by the selection differentail 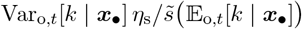, which is the difference between expected knowledge of surviving and all offspring within the lineage. This selection differential is particularly pronounced when there is a significant knowledge difference between offspring (i.e., Var_o,*t*_[*k* | ***x***_•_] is great) and knowledge strongly increases survival (i.e., *η*_s_ is great). The selection differential diminishes as relative differences in survival within the lineage become less pronounced with increasing values of 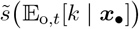. The response of an individual’s expected knowledge to cultural selection due to variation in survival depends on how knowledge is transmitted to the new offspring cohort. It equals 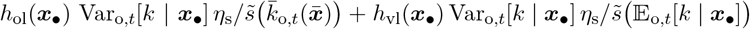 (obtained by substituting the expression of 𝔼_a,*t*+1_[*k* | ***x***_•_] from eq. (10) and 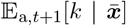 from eq. (10) with 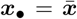 into eq. (9) with *t* = *t* + 1 and identifying the terms corresponding to cultural selection).

Taken together, eqs. (9) and (10) allow us to characterize the expected knowledge at equilibrium. We focus on the expected knowledge in adults only, as this is sufficient to uncover the mechanisms shaping expected knowledge. We show in Appendix A.4.1 that the equilibrium expected adult knowledge in the ***x***_•_-lineage satisfies

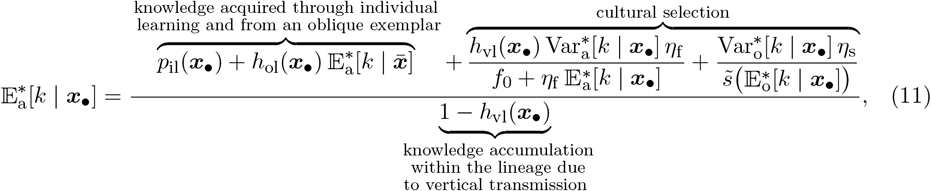

where the superscript ^∗^ indicates that the quantities are evaluated at equilibrium (see also fig. 1d). The first two terms in the numerator of eq. (11) are the expected amount of knowledge acquired through individual and from an oblique exemplar. The remaining terms in the numerator are the increase in expected knowledge driven by cultural selection resulting from differences in fecundity and survival within the lineage due to variation in knowledge. The denominator of eq. (11) captures the inter-generational accumulation of knowledge within the lineage due to vertical transmission. This can be seen by expressing 1*/* 1 − *h*_vl_(***x***_•_) as 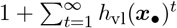, where *h*_vl_(***x***_•_)^*t*^ is the proportion of knowledge acquired by an ancestor *t* generations ago that is effectively transmitted to the focal individual. Since *h*_vl_(***x***_•_) *<* 1, the cumulative contribution of ancestors remains bounded.

We find that offspring exhibit a slightly lower expected knowledge than adults (i.e., 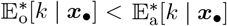, see fig. S.2) because offspring with above-average knowledge are more likely to survive and become adults. In contrast, offspring and adults have the same expected knowledge (i.e., 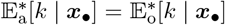, see fig. S.2) when knowledge does not affect survival to adulthood (i.e., *η*_s_ = 0).

#### 3.1.2 Learning stochasticity enhances knowledge by increasing knowledge variance

Our results reveal that cultural selection promotes knowledge accumulation. Because the strength of cultural selection increases with knowledge variance (see eq. (11)), we next investigate the mechanisms shaping this variance. We focus on variance in adults knowledge only, as this is sufficient to identify the mechanisms shaping variance in knowledge. In Appendix A.4.2 we show that the equilibrium variance in the knowledge held by an adult from the ***x***_•_-lineage satisfies

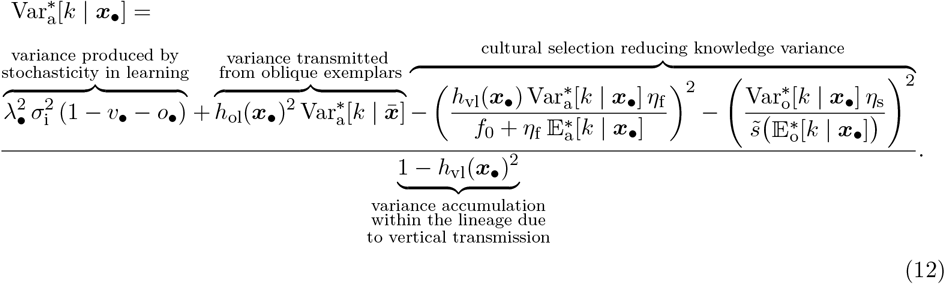

The first term in the numerator of eq. (12) is the knowledge variance produced by stochasticity in individual learning. This term increases with the level of learning stochasticity *σ*_i_, investment in learning *λ*_•_, and the time allocated to individual learning 1 − *v*_•_ − *o*_•_. If learning were deterministic (i.e., *σ*_i_ = 0), no knowledge variance would be produced, and the only solution to eq. (12) would be 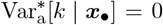 (see proof in Appendix A.4.3). The second term in the numerator accounts for variance transmitted from oblique exemplars. The remaining terms in the numerator are the effect of cultural selection, which reduces knowledge variance. Because survival to adulthood reduces knowledge variance, the equilibrium variance in the knowledge held by an adult 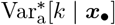 is generally slightly lower than that of an offspring 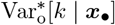 (see fig. S.2). When knowledge does not affect survival (i.e., *η*_s_ = 0), the two variances are equal (i.e., 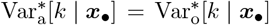, see fig. S.2). Finally, the denominator of eq. (12) describes the accumulation of knowledge variance within the ***x***_•_-lineage through vertical transmission. This can be seen by expressing 1*/*(1 − *h*_vl_(***x***_•_)^2^) as 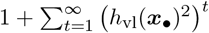, where *h*_vl_(***x***_•_)^2^ is the proportion of knowledge variance from lineage members *t* generations ago that is effectively transmitted to current lineage members.

Altogether eqs. (11) and (12) reveal that, by generating knowledge variance, stochastic individual learning amplifies cultural selection and enhances cumulative knowledge. By numerically estimating 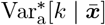 and 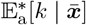 (see Appendix A.4.4 for details on the procedure) for different values of *σ*_i_, we confirm that increased stochasticity in individual learning *σ*_i_ increases population knowledge variance 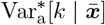, which in turn lead to an increase in mean knowledge 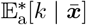 (fig. 2a).

**Figure 2:**
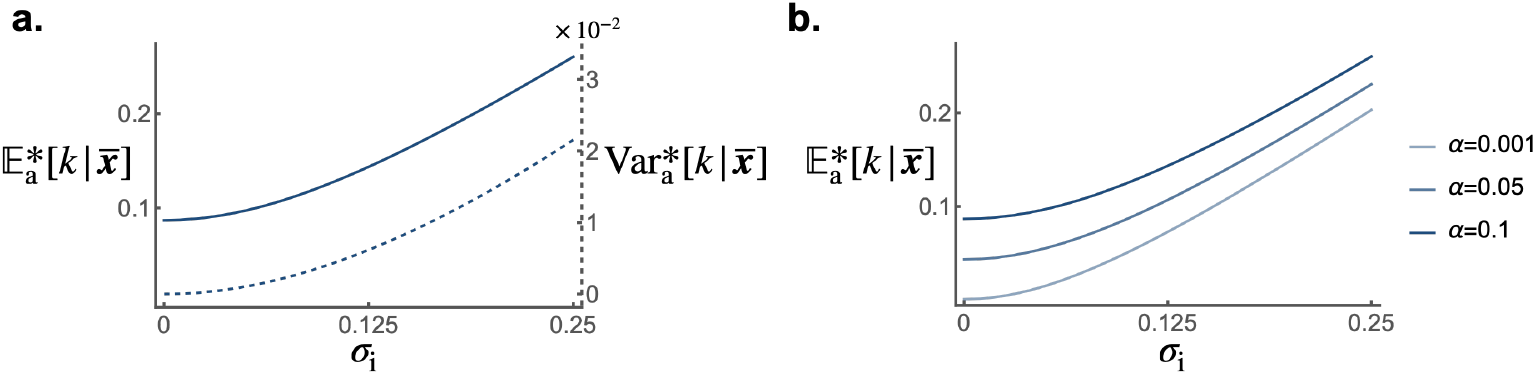
The accumulation of knowledge. **a** Population mean knowledge 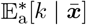 (solid line) and population knowledge variance 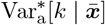 (dashed line) at cultural equilibrium according to intensity of stochastic effects in individual learning *σ*_i_ (left axis gives the scale of cumulative knowledge, and right axis gives the scale of knowledge variance). **b** Population mean knowledge 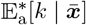 according to intensity of stochastic effects in individual learning *σ*_i_ for different individual learning rates per fraction of investment allocated to learning *α*. Default parameters are: *f*_0_ = 5, *s*_0_ = 1, *β*_v_ = 1.4, *β*_o_ = 1.3, *α* = 0.1, *ϵ* = 0.05, *ρ* = 0.05, *σ*_i_ = 0.1, *η*_f_ = 25, *η*_s_ = 5, ō = 0.2, 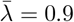.

Remarkably, stochasticity in individual learning *σ*_i_, can foster knowledge accumulation even in populations with low average knowledge-production ability (e.g., *α* = 0.001; see fig. 2b). When stochasticity is particularly high, cultural selection becomes the dominant force driving knowledge acquisition. As a result, populations with different average knowledge-production abilities tend to carry similar levels of mean knowledge, whereas, in the absence of stochasticity, they exhibit marked differences in mean knowledge (see fig. 2b). Additionally, because knowledge improves both fecundity and survival, greater stochasticity in individual learning *σ*_i_ is associated with larger population size (see fig. S.3). The effect of cultural selection in enhancing the mean knowledge 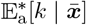 is stronger when knowledge strongly increases fecundity and survival (i.e., when *η*_f_ and *η*_s_ are higher; see fig. S.4), and when vertical and oblique cultural heritability are higher (fig. S.5).

### 3.2 Evolutionary dynamics

#### 3.2.1 Trade-off between fecundity and learning

With knowledge distribution at equilibrium characterized, we now use the selection gradient to investigate the effect of selection on the evolution of the learning traits. By substituting eq. (5) into eq. (6), we obtain the direction of selection on the traits

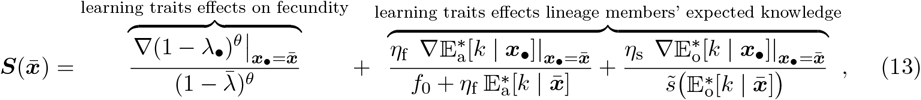

where the gradient operator ∇ is defined in eq. (7).

The first term in eq. (13) describes selection arising from the fecundity costs associated with investment in learning. The remaining terms describe the effect of selection resulting from the impact of learning traits on the expected knowledge of members of a lineage. The strength of these selection pressures diminishes with increasing values of 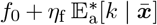 and 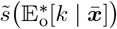, as relative differences in fecundity and survival among lineages become less pronounced.

Overall, eq. (13) indicates that the evolution of learning traits results from a trade-off between the fecundity costs incurred and the knowledge gained. We focus on evaluating the trait values 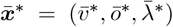 favored by selection in the long-term starting from a population initially expressing 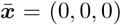 (see Appendix B.3 for details on the procedure). Numerical estimates of the average learning traits at 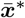 reveal that the fecundity costs can prevent the emergence of learning, particularly when individual learning is inefficient at producing knowledge (i.e., low *α*; see fig. S.6a). As *α* increases, learning can emerge abruptly rather than gradually, because the emergence of learning allows knowledge to accumulate across generations, promoting further social learning and further learning investment in turn.

Fecundity costs also shape the evolution of the learning traits: when the fecundity costs associated with learning investment are higher (i.e., *θ* is higher), individuals invest less in learning (i.e., 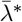 is lower; fig. S.6b). This rapidly limits the evolution of both types of social learning (i.e., 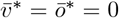 fig. S.6b). Indeed, when the fecundity cost exponent *θ* is higher, individuals allocate more resources to reproduction and less to learning at 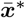 (i.e 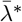 is lower; fig. S.6b), which restricts knowledge production and consequently reduces the benefits of social learning. In the following numerical analyses, we set *θ* = 0.1, a value that permits the evolution of social learning.

#### 3.2.2 Trade-off between different types of learning

To better understand the effect of selection on the learning traits, we decompose their effect on the expected knowledge of adults in the ***x***_•_-lineage. We show in Appendix B.2 that 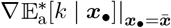 satisfies

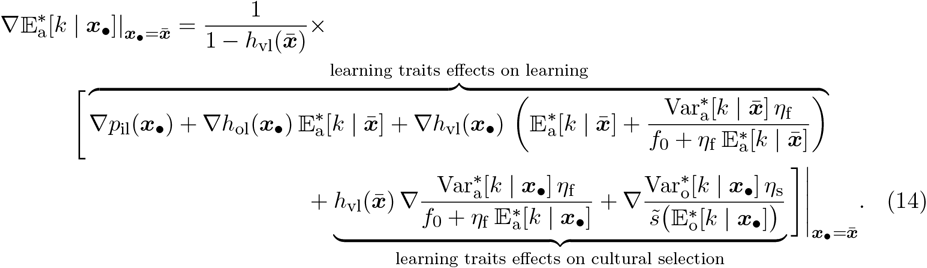

(Appendix B.2). The terms on the second line describe the impact of learning traits on knowledge acquisition. This marginal effect depends on how traits *v, o*, and *λ* influence the knowledge produced individually, as well as that acquired from oblique and vertical exemplars. The marginal knowledge gained through individual learning is 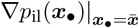. The marginal knowledge gained from oblique and vertical exemplars depends on the effect of the learning traits on the oblique 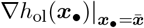 and vertical 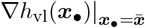 cultural heritability, multiplied by the expected knowledge of a parent and an oblique exemplar. Since the traits *v* and *o* enhance the time allocated to vertical and oblique learning, but reduce the time available for individual knowledge production, their evolution is shaped by trade-offs between different types of learning. The terms on the third line describe the impact of learning traits on cultural selection acting on the expected adult knowledge in the lineage. Learning traits can impact cultural selection by impacting the variance in potential knowledge, as well as the expected fecundity and survival within the lineage.

The selection pressure scales as 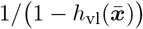, which captures the inter-generational accumulation of the effects of learning traits on knowledge acquisition and on cultural selection, due to vertical transmission. This can be seen by expressing this factor as 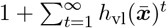, where 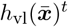 is the population mean proportion of knowledge initially acquired by an ancestor that is transmitted to a direct descendant *t* generations later. This formulation highlights that a change in knowledge in one generation, driven by a change in learning traits, influences the knowledge of all future descendants within the lineage. By capturing the cumulative effects of these interactions, which occur among relatives, the evolutionary dynamics thus incorporate multi-generational kin selection effects.

To understand selection on the learning traits, we also need to examine 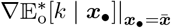 (see eq. (13)). The condition satisfied by 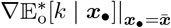 is identical to eq. (14) with 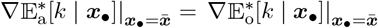 on the left-hand side, minus the effect of learning traits on cultural selection due to differences in survival (see eq. (B.25)). This difference arises because offspring have not yet undergone survival themselves.

Altogether eqs. (13) and (14) reveal that the evolution of learning traits is influenced by the trade-off between knowledge acquisition from different learning types. Numerically estimating the mean trait values 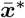 favored by selection reveals that individuals allocate significant time to both vertical and oblique learning (i.e., high 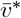 and *ō*^∗^; see fig. S.6c) when knowledge remains relevant over time (i.e., low *ϵ*; e.g., in a stable environment). Under these conditions, individuals can acquire substantial knowledge socially. Since social learning becomes more efficient in stable environments, individuals tend to invest significant resources into learning (i.e., high 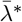 see fig. S.6c). The trade-off between vertical and oblique learning depends on the overlap in knowledge between two adults *ρ*. When there is a high knowledge overlap between adults (i.e., high *ρ*), individuals tend to skip the oblique learning phase at 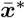 (i.e., *ō*^∗^ = 0; see fig. S.6d), since we assume individuals learn more easily from parent (i.e., *β*_v_ *> β*_o_). Conversely, when there is a low knowledge overlap between adults (i.e., low *ρ*), at 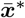 individuals allocate time to oblique learning (i.e., *ō*^∗^ *>* 0), enabling them to acquire knowledge not available from their parents (see fig. S.6d). This outcome is consistent with the results of Maisonneuve et al. (2025) (see AppendixB.4 for details). Although it strongly influences the learning schedule, the overlap in knowledge between two adults *ρ* has little effect on resource allocation between learning and fecundity at the reached evolutionary equilibrium (i.e., little effect on 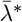 see fig. S.6d).

#### 3.2.3 Learning stochasticity promotes social learning

We now investigate the impact of stochasticity in individual learning, quantified by the parameter *σ*_i_, on the evolution of learning traits. We show that as *σ*_i_ increases, individuals tend to engage more in both vertical and oblique learning at 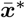 (i.e., 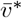 and *ō*^∗^ increase; see fig. 3a). This is because high *σ*_i_ allows the population to accumulate substantial knowledge (see fig. 3g), resulting in cultural exemplars possessing greater knowledge. Under high *σ*_i_, the increased efficiency of social learning then drives individuals to invest more resources in learning at 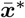 (i.e., high 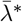 see fig. 3d). This effect is especially pronounced in populations with low knowledge-production ability: stochasticity in learning can drive the evolution of great investment in learning (e.g., 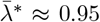 when *α* = 0.025 and *σ*_i_ = 0.25; see fig. S.7), whereas in its absence, investment in learning would be low (e.g., 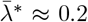 when *α* = 0.025 and *σ*_i_ = 0; see fig. S.7). However, in populations with very low knowledge-production ability, individuals evolve to allocate all resources to fecundity and none to learning, so greater stochasticity in learning has no effect (e.g., *α* = 0.001; see fig. S.7). The impact of *σ*_i_ on learning traits at 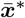 also weakens when the obsolescence rate is high, as rapid knowledge decay across generations limits knowledge accumulation (e.g., *ϵ* = 0.25; fig. S.6e). Although higher *σ*_i_ leads to investing more in learning at the expense of fecundity (i.e., higher 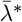 ), the resulting enhancement in population mean knowledge still leads to larger population sizes at 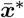 (see fig. 3g).

**Figure 3:**
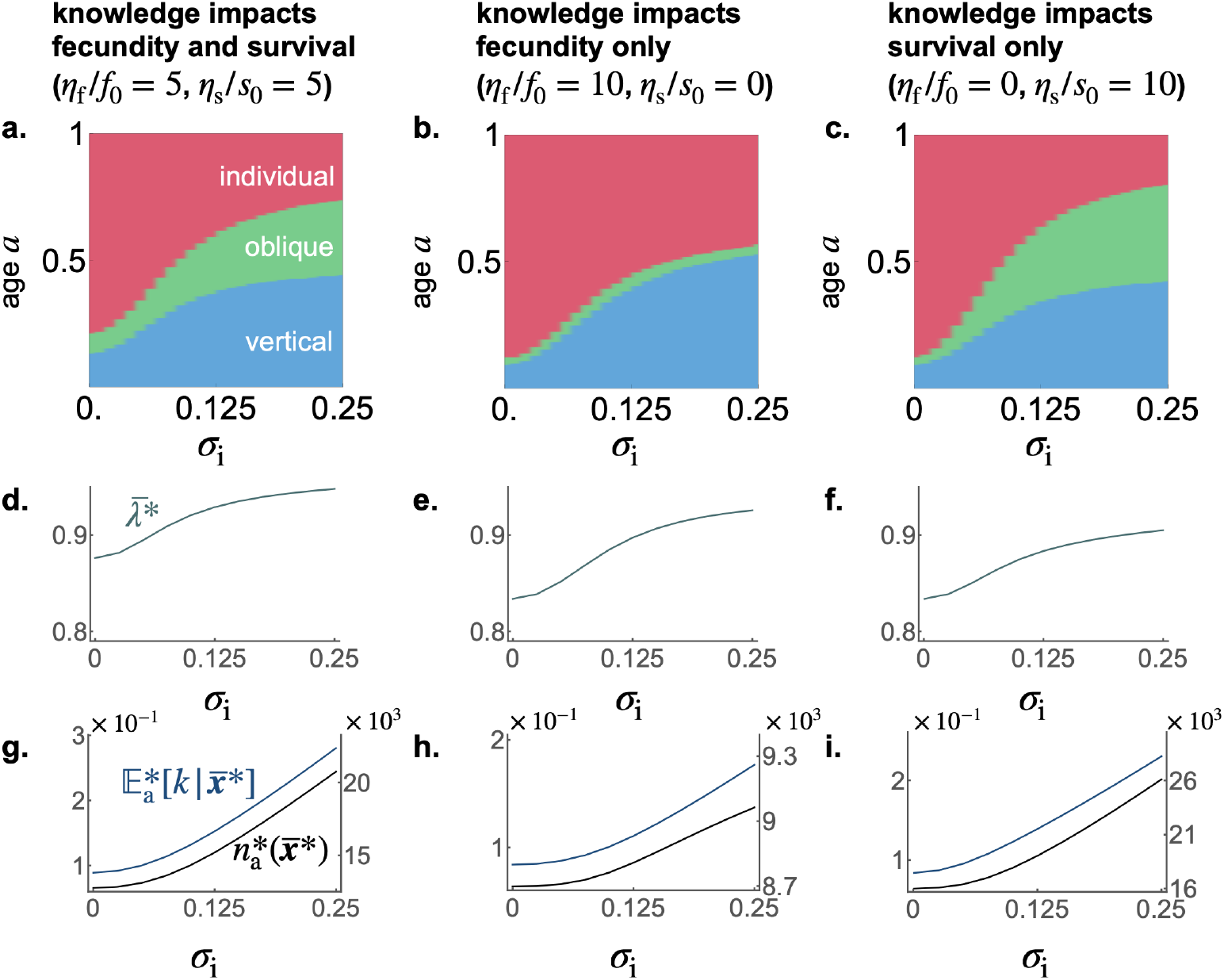
Evolution of learning traits. **a-c** Learning schedule (y-axis) at 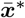 against the intensity of stochastic effects in individual learning *σ*_i_ (x-axis) for different values of the conversion factor that translates knowledge into fecundity *η*_f_ and survival benefits *η*_s_. Blue, green, and pink areas represent time spent performing vertical, oblique, and individual learning, respectively. **d-f** Investment in learning 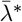 at 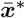 corresponding to panels a-c. **g-j** Population mean knowledge 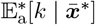 (blue) and adult population size 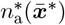 (black) at 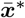 corresponding to panels a-c (left axis gives scale of knowledge, and right axis gives scale of population size, with *γ* = 10^−4^). Default parameters are the same as in fig. 2 with *θ* = 0.1.

Individual-based simulations, which notably relax the assumption of a Gaussian probability density of knowledge, confirm that greater stochasticity in individual learning promotes both increased time allocated to social learning and greater investment in learning at 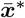 and leads to higher population mean knowledge and population size (see fig. S.9). In addition, simulations that relax the assumption of non-stochastic social learning (i.e., allowing *σ*_v_ *>* 0 or *σ*_o_ *>* 0) reveal similar patterns under increased stochasticity in vertical and oblique learning (see fig. S.10).

However, when adults knowledge significantly overlaps, an increase in *σ*_i_ results in a greater allocation of time to vertical learning and a reduced allocation to oblique learning at 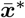 (e.g., increase in 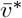 and decrease in *ō*^∗^ when *ρ* = 0.4; see fig. S.6f). In this regime, both parents and other adults provide access to similar knowledge units, so selection favors the most effective social learning mode for acquiring this shared knowledge. Under higher stochasticity in individual learning, the average knowledge of parents 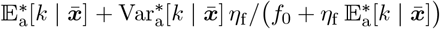 increases more than that of oblique exemplars 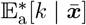 (see fig. S.8) since parents tend to have higher fertility and, consequently, above-average knowledge. As a result, when learning is more stochastic, offspring acquire more of the shared knowledge from their parents.

#### 3.2.4 Selection promotes learning from selected individuals

We now turn to the question of whom individuals should learn from when knowledge varies across the population. Specifically, we investigate which social learning exemplars are favored by natural selection when knowledge improves survival and/or fecundity.

When knowledge enhances fecundity but does not affect survival (i.e., when *η*_f_ *>* 0 and *η*_s_ = 0), increased stochasticity in individual learning (i.e., increased *σ*_i_) leads to allocating more time to vertical learning rather than oblique learning at 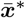 (i.e., a rise in 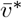 but not in *ō*^∗^; see fig. 3b). In that case, stochasticity in individual learning leads to uncertainty in the knowledge held by potential social learning exemplars. Being a parent then acts as a cue for high fecundity and, therefore, greater knowledge. By contrast, when knowledge increases survival but does not affect fecundity (i.e., when *η*_f_ = 0 and *η*_s_ *>* 0), increased stochasticity in individual learning (i.e., increased *σ*_i_) leads to allocating more time to both vertical and oblique learning at 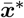 (i.e., a rise in 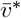 and *ō*^∗^; see fig. 3c). It follows that because adults who are available as social learning exemplars, whether during vertical or oblique learning, have survived until adulthood, their survival is a cue for sizable knowledge. In summary, when stochasticity in learning renders the knowledge of social learning exemplars unpredictable, natural selection favors learning from individuals who exhibit cues for possessing sizable knowledge (e.g., being a parent or having survived to adulthood).

#### 3.2.5 Selection promotes stochasticity in learning

The results so far show that stochasticity in individual learning, quantified by *σ*_i_, markedly affects both the cultural and evolutionary dynamics, and in perhaps counterintuitive ways. Yet, these results may not have traction if *σ*_i_ itself is selected away. Here, we show that learning stochasticity can in fact be favored by selection. We relax the assumption of fixed *σ*_i_ by introducing an additional evolving trait *ζ* ∈ [0, +∞) that increases stochasticity (i.e., 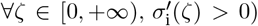. For instance, *ζ* may be a behavioral tendency toward exploration during learning. We assume that learning is inherently stochastic, such that even in the absence of any trait promoting stochasticity (i.e., when *ζ* = 0), a small baseline level of stochasticity remains (i.e., *σ*_*i*_(0) *>* 0, but small). In Appendix C, we show that selection always favors the emergence of the trait *ζ*. This is because in a lineage with traits ***x***_•_ = (*v*_•_, *o*_•_, *λ*_•_, *ζ*_•_), *ζ*_•_ increases knowledge variance (see eq. (C.13)), thereby strengthening cultural selection and leading to higher expected knowledge (see eqs. (C.6) and (C.7)). This effect is further amplified under strong vertical transmission, as increases in expected knowledge driven by cultural selection accumulate across generations within the lineage (see eqs. (C.6) and (C.7)). These results thus confirm the potent role of *σ*_i_ on both the cultural and evolutionary dynamics.

## 4 Discussion

We here first derived the dynamics of individual knowledge in a population using a stochastic model of learning that links generations through vertical and oblique transmission. We showed that as long as the learning process has an element of stochasticity, generating variability in knowledge, and knowledge is transmitted across generations, cultural selection, driven by differences in transmission linked to individual knowledge levels, will inevitably shape the dynamics of knowledge. By estimating the population mean of individual knowledge at equilibrium, our model shows that such cultural selection enhances cumulative knowledge. Those who, by chance, acquire greater knowledge are more likely to survive to adulthood, allowing them to interact with potential learners from the next generation, and to produce more offspring, thereby increasing opportunities for vertical transmission. As a result, individuals with higher knowledge are more likely to pass it on, thereby promoting knowledge accumulation across generations. Our results show that stochasticity in learning amplifies the action of natural selection by promoting variation in knowledge levels.

Our findings are consistent with previous theoretical work in which cultural selection arises because learners preferentially choose more knowledgeable exemplars (e.g., Henrich, 2004; Powell et al., 2009; Kobayashi and Aoki, 2012). In addition, our results highlight that cultural selection can operate even when learners cannot directly assess exemplar knowledge, as long as individuals with greater knowledge are still more likely to transmit, an effect also shown by Cavalli-Sforza and Feldman (1981). Thus, cultural selection is likely to contribute to the improvement of any form of adaptive knowledge that enhances fecundity or survival, even when it is difficult to identify which individuals possess greater knowledge. This may be particularly relevant for knowledge whose benefits are not directly observable because they may be delayed, probabilistic, or rely on hidden causal mechanisms, such as knowledge related to food processing, medicine, rituals, ecological knowledge, or institutional knowledge.

Stochasticity in learning has been extensively studied in reinforcement learning through the effects of random exploration of novel ‘actions’ (which can be thought of as cultural variants in our framework) on learning (Kaelbling et al., 1996; Ladosz et al., 2022; Hao et al., 2024). By increasing the chance of discovering high-reward actions, random exploration enhances the efficiency of selecting and retaining such actions, thereby improving reinforcement learning. While this role of stochasticity is well established, our results demonstrate that it can also enhance the efficiency of social learning, provided knowledge-biased transmission. In stationary environments, the advantages of random exploration in reinforcement learning generally decline as learners approach the optimal action (e.g., Tokic, 2010; Zhang et al., 2023). In contrast, in our model, greater stochasticity remains consistently advantageous because knowledge is unbounded. If we were instead to consider a cultural trait with an optimum (e.g., an ideal tool design or behavior suited to a specific ecological challenge), then, much like in reinforcement learning, stochasticity would likely promote early cultural improvement by enabling the discovery of more adaptive variants, but slow convergence as the population’s traits approach their optimal value. For example, combining individual learning through trial-and-error with social transmission, trait variability across trials can speed convergence when the cultural trait is simple, but increases the distance from the optimum at equilibrium (Lehmann and Wakano, 2013).

Our results also show that, by enhancing cumulative knowledge, the cultural selection introduced by stochasticity in learning favors the evolution of extended social learning phases, as individuals can acquire substantial knowledge from exemplars. In addition, since social learning can provide a large amount of knowledge under these conditions, stochasticity in learning also promotes the evolution of more efficient learning, despite associated fecundity costs. This enables individuals to acquire a greater amount of knowledge during the social learning phase. Our findings may contribute to understanding how the mode of knowledge acquisition differs across knowledge types that vary in the extent of stochasticity in their production. For example, the production of opaque knowledge, such as adaptive taboos (Henrich and Henrich, 2010), adaptive supernatural beliefs (Lightner and Hagen, 2022), or food preparation methods that trigger complex chemical reactions (Beck, 1992), may be particularly stochastic, as their adaptive value is not readily apparent and it is difficult to intentionally produce knowledge that reliably enhances survival and fecundity. Due to this opacity, such knowledge likely became adaptive through a gradual accumulation of random improvements over time. As a result, individuals can acquire far more of this type of knowledge through social learning than they could by independently discovering it. Our results, therefore, predict that opaque knowledge would be primarily acquired socially. This could be the case for knowledge related to food preparation among the Aka Pygmies. Indeed, interviewed individuals reported that they had acquired all of their knowledge related to the preparation of koko, magnoc, and palm wine through social learning (Hewlett and Cavalli-Sforza, 1986). By contrast, other forms of knowledge, such as basic hunting techniques, navigating terrain, or learning to avoid predators, may rely on more transparent causal relationships. As a result, their acquisition may involve less stochasticity and is more likely to occur through individual learning. Nevertheless, stochasticity in knowledge production is likely to explain only part of the variation in modes of knowledge acquisition across knowledge types. Other factors, including the costs of producing knowledge, the need for motor skill practice, the ease of assessing success, and obsolescence, may also play important roles.

Our results reveal that whether knowledge affects fecundity or survival shapes the evolution of learning traits. For instance, when knowledge enhances only fecundity, natural selection favors individuals who mainly learn from their parents rather than from unrelated adults, consistent with the findings of McElreath and Strimling (2008). This is because, when stochasticity introduces variability in knowledge levels, parents tend to have above-average reproductive success and are therefore more likely to possess above-average knowledge. In contrast, randomly chosen adults tend to have average reproductive success and are not particularly likely to possess higher levels of knowledge. Therefore, individuals can acquire more knowledge by learning from their parents than from random adults. One interpretation is that parenthood serves as a cue for possessing knowledge that enhances fecundity, whereas simply being an adult provides no such indication. These findings suggest that knowledge enhancing fecundity should be preferentially acquired from parents. This pattern is supported by empirical evidence from the Aka Pygmies: interviewed individuals reported acquiring, on average, 85.6% of their knowledge about infant care from their parents, compared to 80.7% for knowledge across all domains (Hewlett and Cavalli-Sforza, 1986).

Our results rely on several assumptions that merit further discussion. In particular, we assume that knowledge affects survival and fecundity linearly. This assumption does not affect our conclusions regarding the impact of stochasticity in learning on cumulative knowledge and the evolution of learning traits. However, nonlinear fitness effects could alter knowledge dynamics and the evolutionary trajectories of learning traits, thereby quantitatively affecting our results. For instance, if knowledge yields increasing returns on survival and fecundity, individuals who possess above-average knowledge would gain greater advantages as knowledge accumulates in the population. This would amplify knowledge accumulation, since variation in knowledge would more strongly translate into differences in survival to adulthood and fecundity, thereby increasing cultural selection. As potential exemplars would carry a higher level of knowledge, selection would favor greater reliance on social learning. By contrast, if knowledge yields diminishing returns, we would expect the opposite outcome: knowledge accumulation would slow, and the evolutionary pressure for social learning would weaken.

We also make specific assumptions about the learning process. We assume that learning occurs sequentially through vertical, oblique, and individual phases. Although this is a simplification, it aligns with a general developmental pattern: social learning is more common early in life, while individual learning becomes more prevalent with age (Reader and Laland, 2001; Biesmeijer and Seeley, 2005; Noble et al., 2014; Carr et al., 2015). In humans, the sources of social learning also tend to shift, from primarily vertical transmission during childhood, to greater reliance on oblique transmission in later life stages (Hewlett et al., 2011; Demps et al., 2012; Garfield et al., 2016). In addition, we assume that individuals produce knowledge at a constant average rate, a simplification that allows us to focus on intergenerational knowledge accumulation without explicitly modeling the mechanistic processes underlying individual learning (but see Lehmann and Wakano, 2013, for a model with temporal cultural dynamics with mechanistic individual learning). Finally, we assume that the overlap in knowledge between two adults *ρ* is constant. Allowing *ρ* to emerge endogenously from the learning process could affect our results. Increased stochasticity in learning would reduce overlap in knowledge among adults, making oblique exemplars more likely to possess knowledge not held by parents and thereby promoting greater reliance on oblique learning.

Our study highlights the central role of stochasticity in learning in favoring cumulative knowledge and driving the evolution of learning behaviors. Such stochasticity may be widespread, not only because some level is inevitable, but also because our results show that selection favors increased stochasticity, as it increases the expected knowledge of lineage members when vertical transmission is non-negligible. However, fully understanding the evolution of traits underlying stochasticity in learning requires considering potential trade-offs; for example, a greater tendency to explore may be energetically costly or increase risk exposure. Another trade-off arises because exploratory behavior may slow the rate of knowledge production, as random exploration is more likely to generate non-functional outcomes than functional ones. Incorporating this effect would require explicitly modeling the mechanistic processes underlying individual learning. These insights point to promising directions for future research on the evolutionary dynamics of traits impacting knowledge acquisition and their role in shaping cumulative cultural knowledge.

## Acknowledgments

LM would like to thank Arthur Weyna and Cédric Perret for the useful discussions.

## Author Contributions

LM and LL conceived and designed the study. LM developed the codes and performed the analyses. LM wrote the manuscript with contributions from LL.

## Financial Support

This research received no specific grant from any funding agency, commercial or not-for-profit sectors.

## Conflicts of interest

The authors declare no conflicts of interest.

## Research Transparency and Reproducibility

All the codes used in this study are accessible at github.com/Ludovic-Maisonneuve/stochasticity_evolution_learning.

## Data availability

n/a

## AI declaration

LM has used AI tools to identify grammatical and spelling errors and to enhance the overall fluency of the text.

## Appendix A: Cultural dynamics and equilibrium

Here, we begin by highlighting the similarities and differences between our learning model and previous works (section A.1). We then characterize the outcome of the learning process described by eq. (2) in the absence of stochasticity in social learning (section A.2), and derive the recurrence equations for the expected knowledge and knowledge variance within an ***x***_*•*_-lineage using a Gaussian moment closure (section A.3). Next, we characterize the cultural equilibrium and develop a numerical procedure for computing the population mean knowledge and knowledge variance (section A.4).

### A.1 Connection to previous models

Our learning model, described in the main text and formalized in eq. (2), builds on previous work by Kobayashi et al. (2016) and Maisonneuve et al. (2025), sharing some core assumptions while differing in key aspects. Following Kobayashi et al. (2016), we model stochasticity in both individual and social learning using using white noise from stochastic calculus (see eq. (2) in the main text and eq. (15) in the appendix of Kobayashi et al., 2016). As in Maisonneuve et al. (2025), we assume that learning occurs sequentially through vertical, oblique, and individual phases. A key difference between our model and that of Maisonneuve et al. (2025) lies in the structure of oblique learning: while Maisonneuve et al. (2025) allows individuals to learn from all adults, we assume they learn from only one, reflecting the realistic constraint that offspring cannot interact with everyone. Learning from the entire adult population can inflate the available knowledge, especially under stochastic learning, where some individuals, by chance, accumulate significantly more knowledge than others.

### A.2 Knowledge acquisition without stochasticity in social learning

In this section, we derive eq. (8) using stochastic calculus (e.g., Gardiner, 1985), which provides an expression for computing the realized knowledge acquired by a focal offspring at the end of its learning process, under the assumption that social learning occurs without stochasticity (i.e., *σ*_v_ = *σ*_o_ = 0). When we neglect stochasticity during social learning, the realized knowledge of a focal offspring at each age *a* ∈ [0, 1] with traits ***x***_*•*_ = (*v*_*•*_, *o*_*•*_, *λ*_*•*_), who learn from a parent with knowledge *k*_p*•*_ and an oblique exemplar with knowledge *k*_a*•*_, is a realization of the following stochastic differential equation

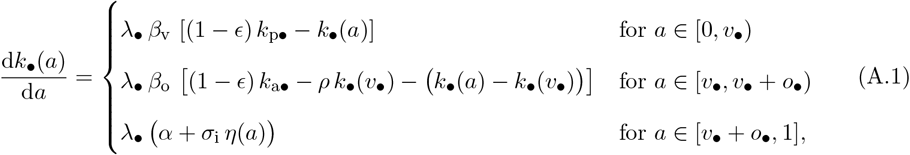

with *k*_*•*_(0) = 0, which is obtained by substituting *σ*_v_ = *σ*_o_ = 0 into eq. (2).

For the final phase of learning, where the dynamics include a stochastic component, eq. (A.1) is interpreted in the Itô sense (Gardiner, 1985, Sec. 4.2, pp. 83–84), whereby

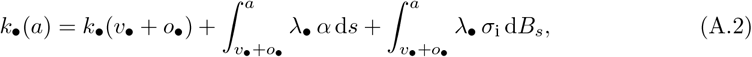

where *B*_*s*_ is the standard Brownian process, which is normally distributed with mean zero and variance *s* (Gardiner, 1985; Bass, 2011, and informally d*B*_*s*_ = *η*(*s*)d*s*).

Accordingly, the knowledge accumulated by the focal offspring at the end of the learning period is

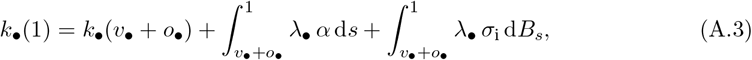

which simplifies to

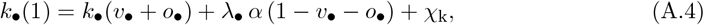

where

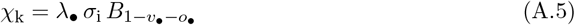

captures the effect of stochastic fluctuations in individual learning. Since 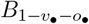 is the realization of a Gaussian random variable with mean 0 and variance 1−*v*_*•*_ −*o*_*•*_, it follows that *χ*_k_ can be obtained as a realization of a Gaussian variable with mean 0 and variance 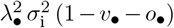.

Substituting the explicit expression for *k*_*•*_(*v*_*•*_ + *o*_*•*_), obtained by solving eq. (A.1) over the deterministic phases, into eq. (A.4) yields

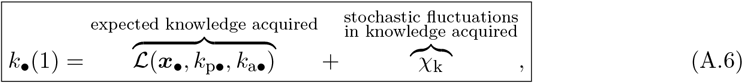

where

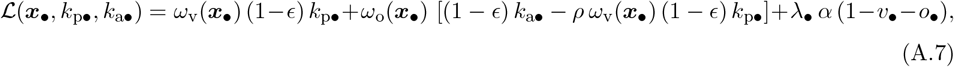

is the expected knowledge acquired. The terms *ω*_v_(***x***_*•*_) and *ω*_o_(***x***_*•*_), given by

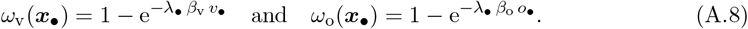

are the proportion of available knowledge at the start of the vertical and oblique learning phases, respectively, that is effectively transmitted to the focal offspring. Substituting the expression for ℒ (***x***_*•*_, *k*_p*•*_, *k*_a*•*_) from eq. (A.7) into eq. (A.6) gives eq. (8) of the main text.

To simplify the notation in what follows, we rewrite eq. (A.7) as

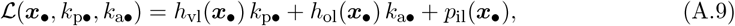

where terms *h*_vl_(***x***_*•*_), *h*_ol_(***x***_*•*_) and *p*_il_(***x***_*•*_) are given by

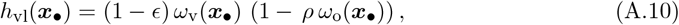

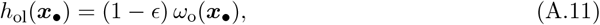

and

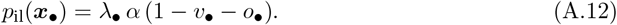

### A.3 Knowledge dynamics

In this section, we derive expressions for the change in the expected knowledge (section A.3.1) and knowledge variance (section A.3.2) in an ***x***_*•*_-lineage. Then we apply a Gaussian closure approximation to derive a closed dynamical system for tracking across generations the expected knowledge and knowledge variance (section A.3.3).

#### A.3.1 Expected knowledge dynamics

Here, we detail how we get eqs. (9) and (10) of the main text, which give the dynamics of the expected knowledge in an ***x***_*•*_-lineage. To this end, it is convenient to define

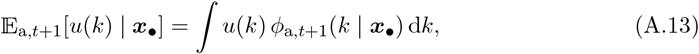

for any function of knowledge *u*(*k*). This quantity denotes the expectation of *u*(*k*) with respect to the probability density *ϕ*_a,*t*+1_(*k* | ***x***_*•*_) that an adult from the ***x***_*•*_-lineage in generation *t* + 1 possesses knowledge *k*. With this notation 𝔼_a,*t*+1_[*k* | ***x***_*•*_] is the expected knowledge of a random adult from the ***x***_*•*_-lineage at generation *t* + 1 (here and throughout the subscript _a,*t*_ denotes quantities computed among adults of generation *t* for all *t* ≥ 1; see fig. A.1).

By definition,

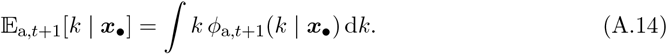

**Figure A.1:**
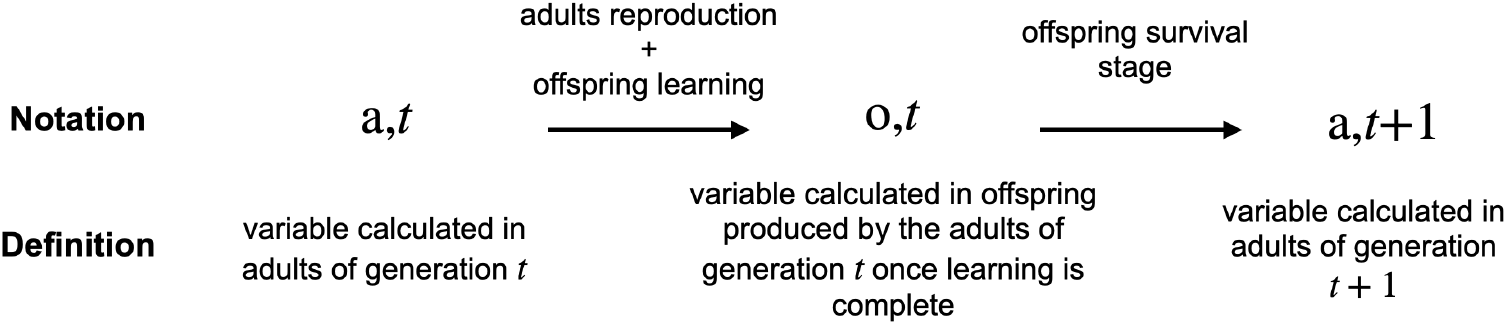
Notation specifying the stage in the life cycle and generation at which variables are calculated.

The probability density that a random adult from the ***x***_*•*_-lineage of generation *t* + 1 possesses knowledge *k, ϕ*_a,*t*+1_(*k* | ***x***_*•*_), can be derived from the probability density that a random offspring produced by an adult from the ***x***_*•*_-lineage of generation *t* possesses knowledge *k, ϕ*_o,*t*_(*k* | ***x***_*•*_) (here and throughout, the subscript _o,*t*_ denotes quantities computed among an offspring produced by an adult of generation *t* after learning is complete and before the survival stage; see fig. A.1), who will potentially become one of the adults of generation *t* + 1. The probability density of knowledge of a random offspring is then weighted by the relative survival probability associated with each possible knowledge:

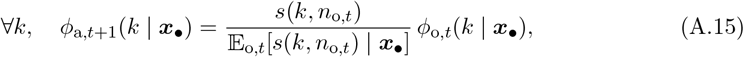

where *n*_o,*t*_ is the number of offspring produced by adults of generation *t* and *s*(*k, n*_o,*t*_) is the survival probability associated with traits ***x***_*•*_ and knowledge *k*. For any function of knowledge *u*(*k*),

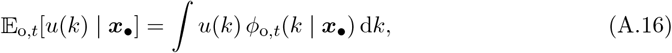

denotes the expectation of this function over the probability density of knowledge *ϕ*_o,*t*_(*k* | ***x***_*•*_) of a random offspring produced by an adult of generation *t*. Here, 𝔼 _o,*t*_[*s*(*k, n*_o,*t*_) | ***x***_*•*_] is the expected survival probability of an offspring produced by an adult of the ***x***_*•*_-lineage of generation *t*. Note that because *ϕ*_a,*t*+1_(*k* | ***x***_*•*_) is a probability density of knowledge among surviving offspring within the ***x***_*•*_-lineage, the survival probability in eq. (A.15) is normalized by the expected survival probability in that lineage 𝔼 _o,*t*_[*s*(*k, n*_o,*t*_) | ***x***_*•*_], not by the expected survival probability in the population.

Noting from eqs. (3) and (4) with *k*_o*•*_ = *k* that *s*(*k, n*_o,*t*_) depends linearly on *k* and using that ∫*ϕ*_o,*t*_(*k* | ***x***_*•*_) d*k* = 1 we have

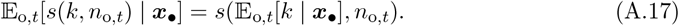

By substituting the expressions of *ϕ*_a,*t*+1_(*k* | ***x***_*•*_) and 𝔼 _o,*t*_[*s*(*k, n*_o,*t*_) | ***x***_*•*_] from eqs. (A.15) and (A.17) into eq. (A.14) we obtain

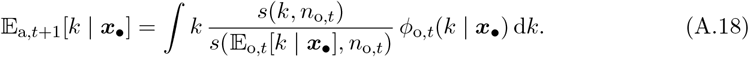

Next, we rewrite *s*(*k, n*_o,*t*_)*/s*(𝔼 _o,*t*_[*k* | ***x***_*•*_], *n*_o,*t*_). Using the expression for *s*(*k, n*_o,*t*_) and *s*(𝔼 _o,*t*_[*k* | ***x***_*•*_], *n*_o,*t*_) from eq. (3) with *k*_o*•*_ = *k, n*_o_ = *n*_o,*t*_ and with *k*_o*•*_ = 𝔼 _o,*t*_[*k* | ***x***_*•*_], *n*_o_ = *n*_o,*t*_ we obtain

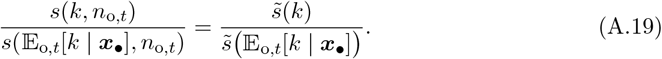

By substituting 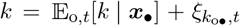, where 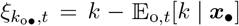, into eq. (A.19) and noting from eq. (4) with *k*_o*•*_ = *k* that 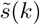 depends linearly on *k* with a linear coefficient *η*_s_, we obtain

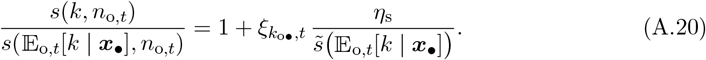

Substituting 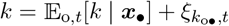 and eq. (A.20) into eq. (A.18) gives

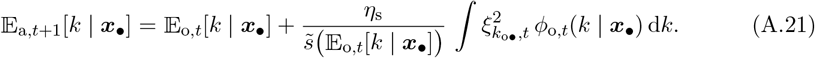

Noting that the integral is the variance in the knowledge held by a random offspring produced by an adult from the ***x***_*•*_-lineage at generation *t*, denoted as Var_o,*t*_[*k* | ***x***_*•*_], we obtain

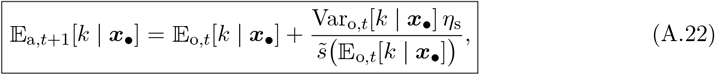

which is eq. (10) from the main text.

Note that, in eq. (A.22), the term 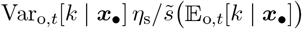 captures the effect of cultural selection arising from differential survival among lineage members, due to variation in knowledge.

To obtain the expected knowledge dynamics across one generation, we need to determine the expected knowledge of a random offspring produced by an adult of generation *t*, 𝔼 _o,*t*_[*k* | ***x***_*•*_]. By definition,

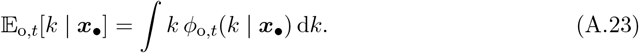

For each offspring, the knowledge acquired depends on the knowledge of both the parent and the oblique exemplar, as well as on the outcome of the stochasticity in the learning process. By integrating over the probability density of parental knowledge, the probability density of knowledge among potential oblique exemplars, and the probability density of stochastic learning outcomes, we obtain

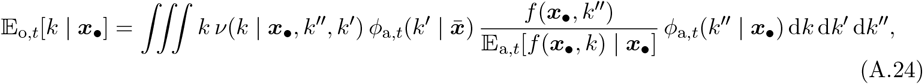

where 𝔼 _a,*t*_[*f* (***x***_*•*_, *k*) | ***x***_*•*_] is the mean fecundity of adults belonging to the ***x***_*•*_-lineage at generation *t* and is equal to

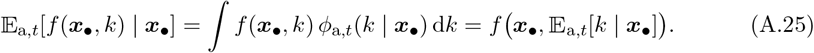

The integral over *k*^*′′*^ encompasses all possible knowledge value of parents, weighted by the probability density *f* (***x***_*•*_, *k*^*′′*^)*/*𝔼_a,*t*_[*f* (***x***_*•*_, *k*) | ***x***_*•*_] *ϕ*_a,*t*_(*k*^*′′*^ | ***x***_*•*_) that a parent has knowledge *k*^*′′*^. Each parent’s contribution depends on its fecundity relative to the lineage’s mean fecundity, *f* (***x***_*•*_, *k*^*′′*^)*/*𝔼_a,*t*_[*f* (***x***_*•*_, *k*) | ***x***_*•*_], because lineage members with higher fecundity are more likely to transmit their traits and knowledge.

The integral over *k*^*′*^ encompasses all possible knowledge values of oblique exemplars in generation *t*, weighted by the probability density 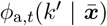 that an oblique exemplar has knowledge *k*^*′*^. Since the variances of population traits are small, we approximate the population as being quasi-monomorphic for the mean traits 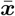, and thus assume that offspring acquire knowledge obliquely from adults with traits 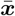.

Finally, the integral over *k* encompasses all possible outcomes of the learning process, weighted by the probability density *ν*(*k* | ***x***_*•*_, *k*^*′′*^, *k*^*′*^) of the offspring’s knowledge *k* at the end of the learning process, given parental knowledge *k*^*′′*^ and oblique learning examplar knowledge *k*^*′*^.

From eq. (A.6) with *k*_*•*_(1) = *k*, we have that *k* = ℒ (***x***_*•*_, *k*^*′′*^, *k*^*′*^) + *χ*_k_ where ℒ (***x***_*•*_, *k*^*′′*^, *k*^*′*^) is deterministic and *χ*_k_ is a realization a Gaussian random variable with mean 0 and variance 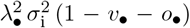. Since *k* is the sum of a deterministic term ℒ (***x***_*•*_, *k*^*′′*^, *k*^*′*^) and a random fluctuation *χ*_k_, its probability density has the same shape as that of *χ*_k_, but centered around ℒ (***x***_*•*_, *k*^*′′*^, *k*^*′*^) instead of 0. This means the probability density of *k* is obtained by shifting the density of *χ*_k_ horizontally so that its mean aligns with ℒ (***x***_*•*_, *k*^*′′*^, *k*^*′*^)

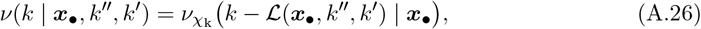

where *νχ*_*k*_ is a Gaussian probability density function with mean 0 and variance 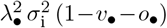. We obtain eq. (A.26) by noting that *k* − ℒ(***x***_*•*_, *k*^*′′*^, *k*^*′*^) = *χ*_k_, so the probability that *k* takes a particular value is the same as the probability that *χ*_k_ takes the value *k* − *L*(***x***_*•*_, *k*^*′′*^, *k*^*′*^).

By substituting the expressions of 𝔼_a,*t*_[*f* (***x***_*•*_, *k*) | ***x***_*•*_] and *ν*(*k* | ***x***_*•*_, *k*^*′′*^, *k*^*′*^) from eqs. (A.25) and (A.26) into eq. (A.24) and by performing a change of variable *χ*_k_ = *k* − ℒ (***x***_*•*_, *k*^*′′*^, *k*^*′*^) we obtain

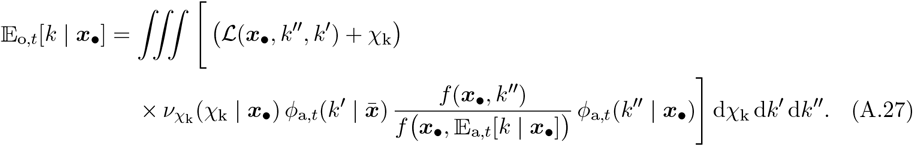

Equation (A.27) can be rearranged as follows

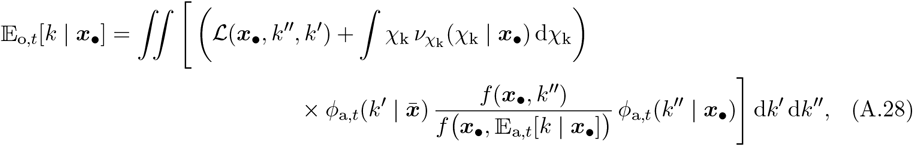

and, because *χ*_k_ is centered at 0 it simplifies to

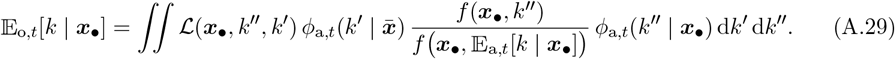

Next, we expand *f* (***x***_*•*_, *k*^*′′*^)*/f* ***x***_*•*_,𝔼_a,*t*_[*k* | ***x***_*•*_] . By substituting 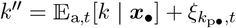, where 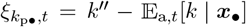, and noting from eq. (1) that *f* (***x***_*•*_, *k*^*′′*^) depends linearly on *k*^*′′*^, we obtain

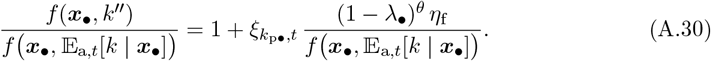

By substituting the expression of *f* ***x***_*•*_, 𝔼_a,*t*_[*k* | ***x***_*•*_] from eq. (1) with *k*^*′′*^ = 𝔼_a,*t*_[*k* | ***x***_*•*_] into the right-hand side of eq. (A.30) we obtain

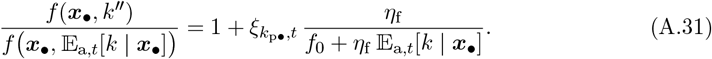

We now rewrite ℒ (***x***_*•*_, *k*^*′′*^, *k*^*′*^). Because ℒ (***x***_*•*_, *k*^*′′*^, *k*^*′*^) is linear on *k*^*′′*^ and *k*^*′*^ with coefficient of linearity *h*_vl_(***x***_*•*_) and *h*_ol_(***x***_*•*_) (see eq. (A.9)), by substituting 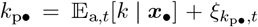 and 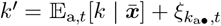 where 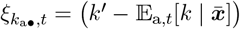, we obtain

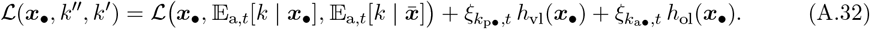

Substituting eqs. (A.31) and (A.32) into eq. (A.29), and noting that 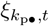 and 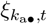 are centered around zero and are independent, we obtain

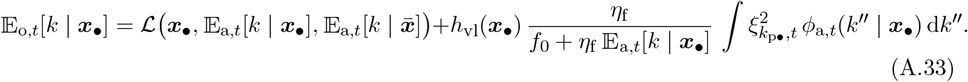

Noting that the integral is the variance in the knowledge held by a random adult from the ***x***_*•*_-lineage at generation *t*, denoted as Var_a,*t*_[*k* | ***x***_*•*_], we obtain

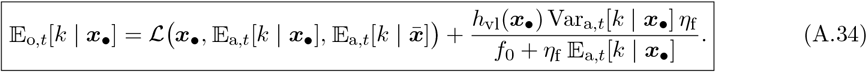

By substituting the expression of 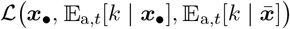 from eq. (A.9) with *k*_p*•*_ = 𝔼_a,*t*_[*k* | ***x***_*•*_] and 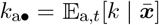 gives eq. (9) from the main text.

Note that, in eq. (A.34), the term *h*_vl_(***x***_*•*_) Var_a,*t*_[*k* | ***x***_*•*_] *η*_f_ */( f*_0_ + *η*_f_ 𝔼_a,*t*_[*k* | ***x***_*•*_]) captures the effect of cultural selection arising from differential fecundity among lineage members, due to variation in knowledge.

Equations (A.22) and (A.34) give the dynamics of the expected knowledge within the ***x***_*•*_-lineage.

#### A.3.2 Knowledge variance dynamics

Equations (A.22) and (A.34) enable us to track changes in expected knowledge within a lineage across a single generation. However, to iterate these equations also requires tracking the variance in the knowledge held by a lineage member simultaneously. In this section, we derive the expression of the dynamics of the variance in the knowledge held by a random adult within the ***x***_*•*_-lineage.

By definition, the variance in the knowledge held by a random adult from the ***x***_*•*_-lineage at generation *t* + 1, Var_a,*t*+1_[*k* | ***x***_*•*_], is given by

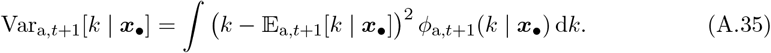

Using 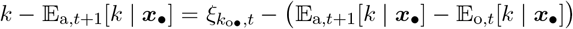 and 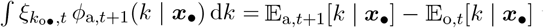 we can expand eq. (A.35) as follows

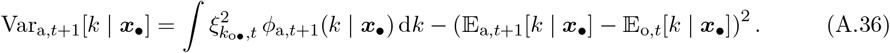

By substituting the expression of *ϕ*_a,*t*+1_(*k* | ***x***_*•*_) from eq. (A.15) into eq. (A.36), we obtain

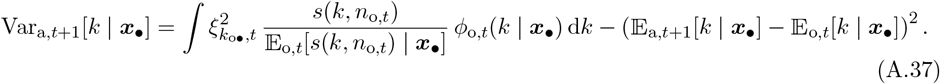

Substituting the expression of *s*(*k, n*_o,*t*_)*/*𝔼_o,*t*_[*s*(*k, n*_o,*t*_) | ***x***_*•*_] from eq. (A.20) (with *s*(𝔼_o,*t*_[*k* | ***x***_*•*_], *n*_o,*t*_) = 𝔼_o,*t*_[*s*(*k, n*_o,*t*_) | ***x***_*•*_]) into eq. (A.37) we find

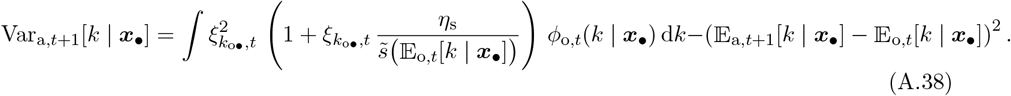

By expanding the integral in eq. (A.38) we find

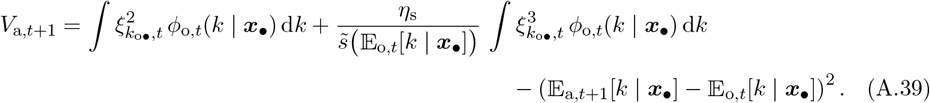

Noting that

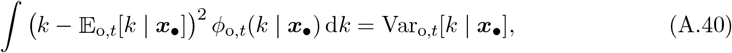

and

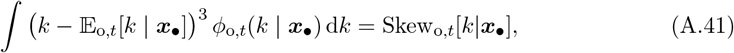

where Skew_o,*t*_[*k*|***x***_*•*_] is the skewness in the knowledge held by a random offspring produced by an adult from the ***x***_*•*_-lineage at generation *t*, and by substituting the expression for 𝔼_a,*t*+1_[*k* | ***x***_*•*_] from eq. (A.22) into eq. (A.39), we obtain

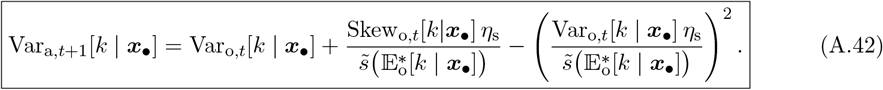

We now compute the variance in the knowledge held by a random offspring produced by an adult from the ***x***_*•*_-lineage at generation *t*, Var_o,*t*_[*k* | ***x***_*•*_]. The variance Var_o,*t*_[*k* | ***x***_*•*_] can be obtained by integrating over the probability density of parental knowledge, the probability density of knowledge among potential oblique exemplars, and the probability density of all stochastic learning outcomes to obtain

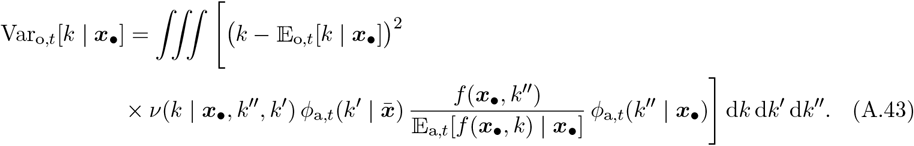

By substituting the expressions of 𝔼_a,*t*_[*f* (***x***_*•*_, *k*) | ***x***_*•*_] and *ν*(*k* | ***x***_*•*_, *k*^*′′*^, *k*^*′*^) from eqs. (A.25) and (A.26) into eq. (A.43) and by performing a change of variable *χ*_k_ = *k* − ℒ (***x***_*•*_, *k*^*′′*^, *k*^*′*^) we obtain

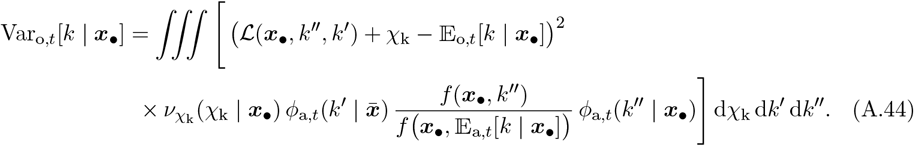

By substituting the expression of *f* (***x***_*•*_, *k*^*′′*^)*/f* ***x***_*•*_, 𝔼_a,*t*_[*k* | ***x***_*•*_] and ℒ (***x***_*•*_, *k*^*′′*^, *k*^*′*^) from eqs. (A.31) and (A.32) into eq. (A.44) we obtain

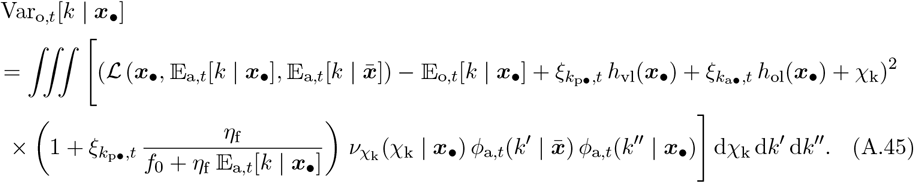

Expanding the square in eq. (A.45), and noting that *χ*_k_ is centered at zero, we get

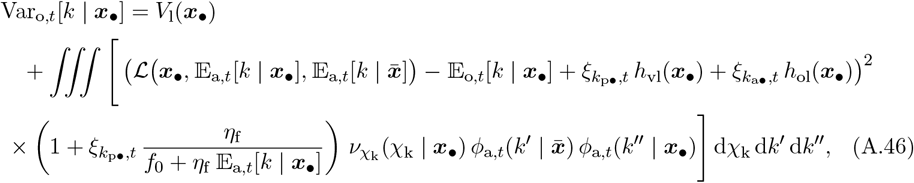

where

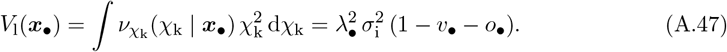

Expanding the integral in eq. (A.46) and noting that 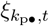 and 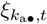 are centered around zero and independent, we find

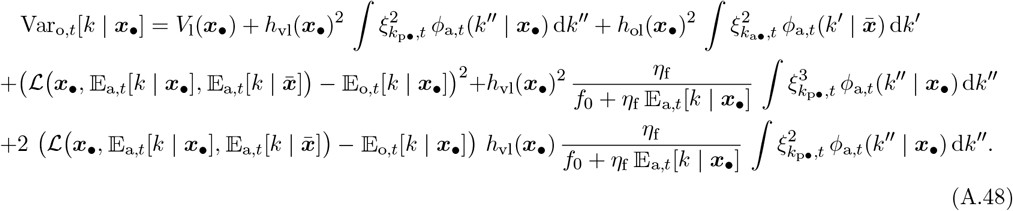

Noting that

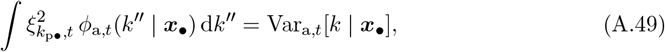

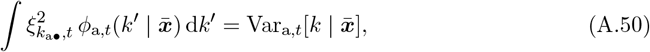

and

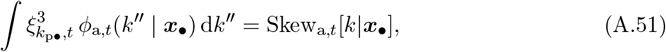

where Skew_a,*t*_[*k*|***x***_*•*_] is the skewness in the knowledge held by a random adult from the ***x***_*•*_-lineage at generation *t* and by substituting the expression for 𝔼_o,*t*_[*k* | ***x***_*•*_] from eq. (A.34) into eq. (A.48) we obtain

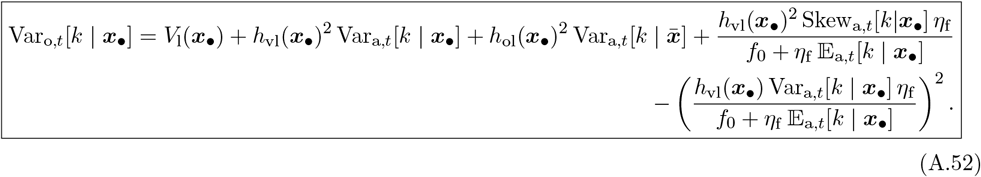

Equations (A.42) and (A.52) give the dynamics of knowledge variance.

#### A.3.3 Gaussian closure approximation

Equations (A.22), (A.34), (A.42), and (A.52) describe the dynamics of the expected knowledge and knowledge variance in a lineage. However, tracking the expected knowledge and knowledge variance requires tracking knowledge skewness, which in turn necessitates tracking fourth-order moments of knowledge probability density. Since the dynamics of each moment depend on higher-order moments, tracking the cultural dynamics entails following the dynamics of the infinite sequence of moments. To circumvent this issue, we apply a Gaussian closure approximation, assuming that the probability density of knowledge within a lineage can be approximated by a Gaussian probability density. This assumption allows us to track the evolution of the probability density of knowledge across generations by tracking only its mean and variance.

Under the Gaussian closure approximation, the knowledge skewness is zero. By substituting Skew_a,*t*_[*k*|***x***_*•*_] = Skew_o,*t*_[*k*|***x***_*•*_] = 0 into the system of equations formed by eqs. (A.22), (A.34), (A.42) and (A.52) we obtain

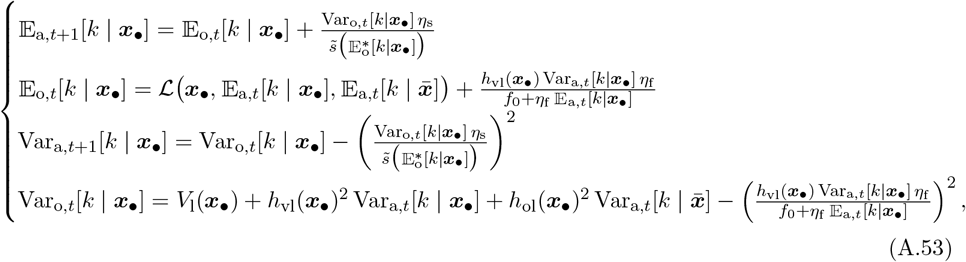

which characterized the dynamics of the expected knowledge and knowledge variance of the ***x***_*•*_-lineage member.

### A.4 Probability density of knowledge at cultural equilibrium

In this section, we first characterize the equilibrium conditions for the expected knowledge and knowledge variance at cultural equilibrium (sections A.4.1 and A.4.2). Next, we demonstrate that in the absence of stochasticity in learning, there is no knowledge variance at the cultural equilibrium (section A.4.3). Finally, we describe the numerical method used to estimate the equilibrium values of mean knowledge and knowledge variance in the population (section A.4.4).

#### A.4.1 Characterization of the expected knowledge at cultural equilibrium

Here, we derive eq. (11) from the main text, which specifies the condition satisfied by the expected adults knowledge 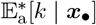 at equilibrium (here and hereafter the notation ^∗^ denotes a variable at equilibrium).

At equilibrium, the expected knowledge and the variance in knowledge for randomly chosen adults and offspring remain unchanged across generations. By substituting 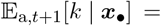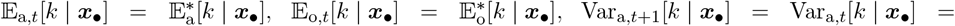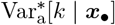 and 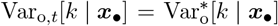 into the dynamical system eq. (A.53), we obtain the following conditions that these equilibrium values must satisfy

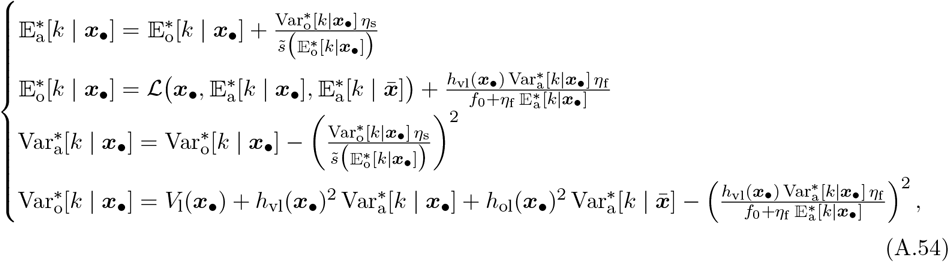

where 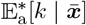 and 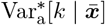 are the population’s mean and variance of adult knowledge, which verifies eq. (A.54) with 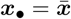.

By replacing 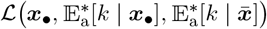 with its expression from eq. (A.9) (with 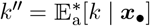 and 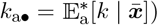 in the second line of eq. (A.54), we obtain

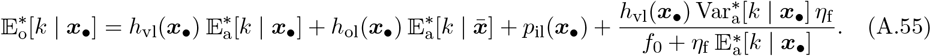

By substituting the expression for 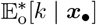 from eq. (A.55) into the first line of eq. (A.54), we obtain

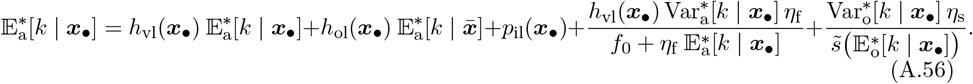

This can be rewritten as

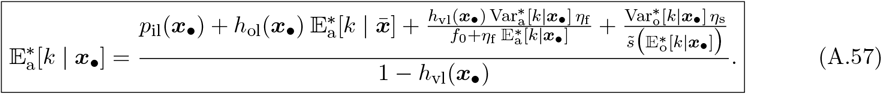

Equation (A.57) is eq. (11) from the main text.

#### A.4.2 Characterization of the knowledge variance at cultural equilibrium

Here, we derive eq. (12) from the main text, which specifies the condition satisfied by the variance in the knowledge held by a random adult 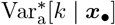 at equilibrium. By substituting the expression for 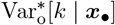 from the fourth line of eq. (A.54) into the third line of the same equation, we obtain

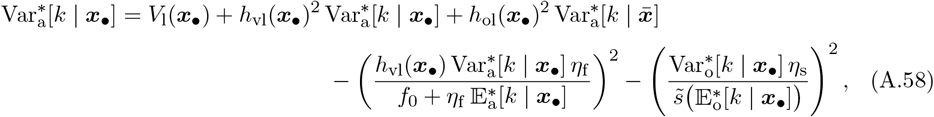

which can be rearranged as

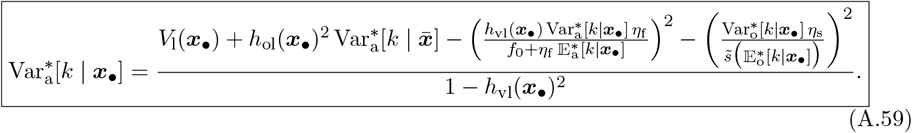

Substituting the expression for *V*_l_(***x***_*•*_) from eq. (A.47) into eq. (A.59) gives eq. (12) from the main text.

#### A.4.3 Knowledge variance in the absence of stochasticity in learning

Here, we show that if individual learning is deterministic (i.e., *σ*_i_ = 0), the equilibrium variance of adult knowledge is zero (i.e., 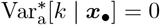).

We assume that, initially, all individuals have no knowledge, so that the knowledge variance is null: 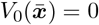. We aim to prove that this remains true for all generations *t*, i.e., 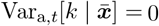 for all *t*. Using mathematical induction, this is equivalent to showing that 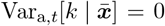 implies 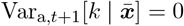.

Assume that 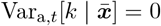. Substituting *σ*_i_ = 0 into eq. (A.47), we obtain *V*_l_(***x***_*•*_) = 0. Then, substituting both 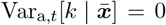 and *V*_l_(***x***_*•*_) = 0 into the fourth line of eq. (A.53), we find that Var_o,*t*_[*k* | ***x***_*•*_] = 0. Finally, substituting Var_o,*t*_[*k* | ***x***_*•*_] = 0 into the third line of eq. (A.53) yields 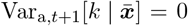. This completes the inductive step and shows that 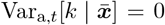 for all *t*, which implies that at equilibrium, 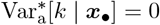. This concludes the proof.

#### A.4.4 Numerical estimation of population mean and variance in knowledge

To numerically determine the mean and variance of knowledge within the population at cultural equilibrium, 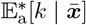 and 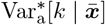, we iterated the dynamics described by eq. (A.53) with 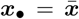 across generations, starting from the initial conditions 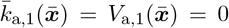. The expressions for 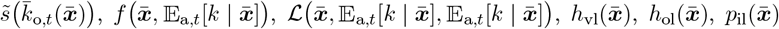 and *V*_l_(***x***_*•*_) are given by eqs. (4) (with 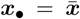 and 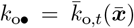), (1) (with 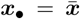 and 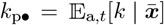), (A.9) (with 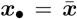 and 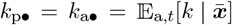), (A.10) (with 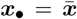), (A.11) (with 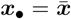), (A.12) (with 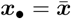) and (A.47) (with 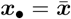) respectively, treating 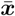 as a parameter. We iterate this dynamics until 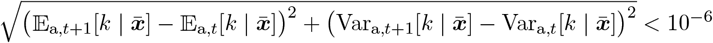.

## Appendix B: Evolutionary dynamics

In Appendix B, we derive eq. (5) from the main text, which provides a simple expression for the lineage fitness (section B.1). We then derive eq. (14) from the main text, which establishes the conditions that 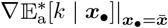 and 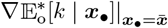 satisfy (section B.2). Lastly, we detail the numerical procedure for computing the values of mean traits favored by selection (section B.3).

### B.1 Lineage fitness

In this section, we detail how to obtain eq. (5) from the main text, which provides an expression for the lineage fitness 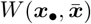 of an ***x***_*•*_-lineage introduced into a population with mean trait values 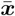. The lineage fitness satisfies

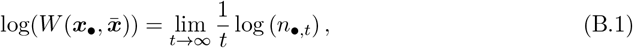

where the sequence (*n*_*•,t*_)_*t*≥0_ is a realization of a stochastic process, where *n*_*•,t*_ describes the random number of individuals in the ***x***_*•*_-lineage at time *t* following the introduction of the ***x***_*•*_-mutant (Cohen, 1979; Caswell, 2000). This number changes according to the recursion

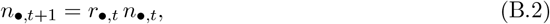

with initial condition *n*_*•*,0_ = 1. Here, *r*_*•,t*_ is the realized growth rate, i.e., the average number of viable offspring produced by individuals in the ***x***_*•*_-lineage, *t* generations after the introduction of the ***x***_*•*_-mutant. This quantity depends on multiple factors, including the knowledge individuals carry, the number of offspring they produce, the learning outcomes of the offspring, and their survival based on their knowledge. Since offspring acquire knowledge from both their parent and randomly selected adults in the population, the probability density of *r*_*•,t*_ depends in part on the distribution of knowledge within the lineage and across the population in the parental generation.

Using eq. (B.2) with *t* = *h*, for *h* ∈ {0, …, *t* − 1}, we obtain

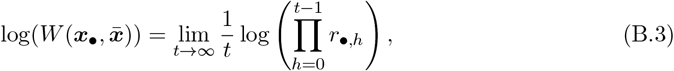

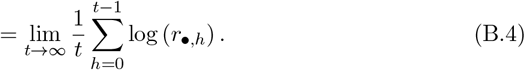

We assume that the stochastic process underlying the lineage process ***x***_*•*_ growth reaches a stationary regime, such that *r*_*•,t*_ converges in distribution to a random variable *r*_*•*_. By applying the ergodic theorem for population growth (Caswell, 2000), eq. (B.3) can then be expressed as

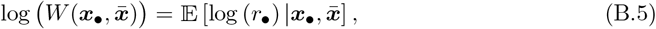

where 𝔼 denotes the expectation under the stationary probability density of *r*_*•*_. It depends on the trait values within the lineage, ***x***_*•*_, as these directly influence learning and fecundity, and also determine the stationary probability density of knowledge within the lineage. It also depends on the population mean trait values 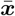, which determine the equilibrium population size and the stationary probability density of knowledge in the population.

The variance in *r*_*•,t*_ stems from generational differences in the knowledge held by members of the lineage. We assume that the variability in knowledge is small, resulting in limited fluctuations in *r*_*•,t*_ at stationarity. Using a Taylor expansion of log (*r*_*•*_) around 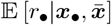, we obtain

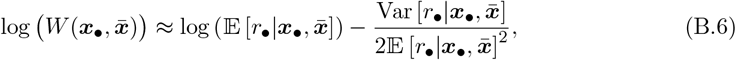

where Var denotes the variance under the stationary probability density of *r*_*•,t*_. By neglecting the term proportional to 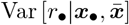, we obtain

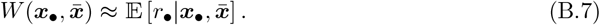

While this is an approximation of lineage fitness, the estimated trait values reached at the evolutionary equilibrium using this approximation (see section B.3 for details) are consistent with those observed in individual-based simulations (see fig. S.9), suggesting that eq. (B.7) provides a reliable expression of lineage fitness.

The expected growth rate of the lineage 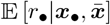 equals the expected number of viable offspring for a random lineage member under the stationary distribution. Accordingly, the lineage fitness 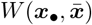 can be written as

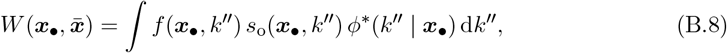

where *ϕ*^∗^(*k*^*′′*^ | ***x***_*•*_) is the stationary probability density that an adult with traits ***x***_*•*_ carries knowledge value *k*^*′′*^ at cultural equilibrium, and the integral is taken over all possible values of *k′′*.

The term *f* (***x***_*•*_, *k*^*′′*^) (whose expression is given in eq. (1)) is the expected number of offspring produced by an adult with traits ***x***_*•*_ and knowledge *k*^*′′*^. The term

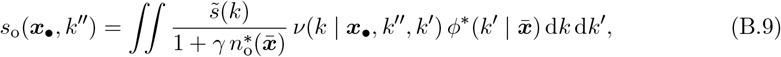

is the expected probability of survival for a random offspring of an adult with traits ***x***_*•*_ and knowledge *k*^*′′*^. The survival probability of a descendant offspring depends on its trait, which is ***x***_*•*_ in the ***x***_*•*_-lineage, and on its knowledge, *k*, which is a random variable. To account for all possible survival outcomes, we must consider the full probability density of possible offspring knowledge values.

Since offspring’s knowledge results from learning from an oblique exemplar, first we integrate over the probability density of the exemplar’s knowledge *k*^*′*^. Given that the variances of population traits are small, we approximate the population as being quasi-monomorphic for the mean traits 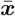. Consequently, we assume that offspring acquire knowledge obliquely from adults with traits 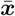, which carry knowledge *k*^*′*^ with probability density 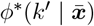. Then, for each possible oblique exemplar knowledge value *k*^*′*^, we integrate over all possible offspring knowledge values *k*, weighted by the probability density *ν*(*k* | ***x***_*•*_, *k*^*′′*^, *k*^*′*^) that an offspring with trait ***x***_*•*_, parental knowledge *k*^*′′*^, and exemplar knowledge *k*^*′*^ acquires knowledge *k*. Since *k* = *k*_*•*_(1), where *k*_*•*_(*a*) is a realization of the stochastic differential equation eq. (2), the probability density *ν*(*k* | ***x***_*•*_, *k*^*′′*^, *k*^*′*^) is entirely determined by eq. (2) (see, for example, the expression of *ν*(*k* | ***x***_*•*_, *k*^*′′*^, *k*^*′*^) given in eq. (A.26) when *σ*_v_ = *σ*_o_ = 0). Finally, 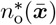 is the expected number of offspring produced at demographic and cultural equilibrium in a population with mean traits 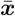.

Using the expression for 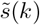 and *s*_o_(***x***_*•*_, *k*^*′′*^) from eqs. (4) and (B.9) with *k*_o*•*_ = *k*, we rewrite eq. (B.8) as

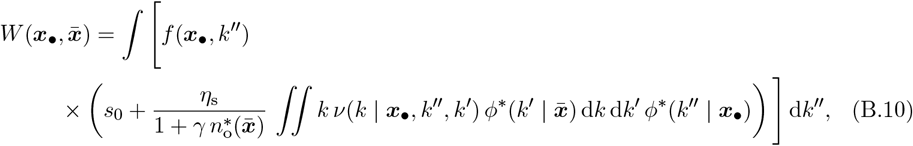

which can be rewritten as

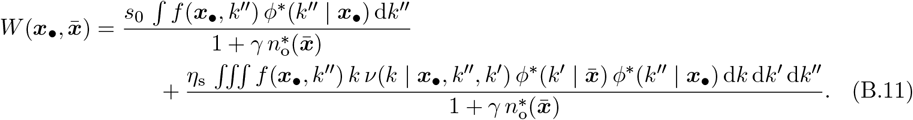

Because *f* (***x***_*•*_, *k*^*′′*^) depends linearly on *k*^*′′*^ and ∫*ϕ*^∗^(*k*^*′′*^ | ***x***_*•*_) d*k*^*′′*^ = 1 we have

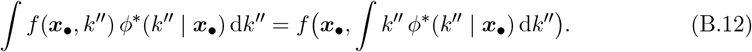

By definition, ∫*k*^*′′*^ *ϕ*^∗^(*k*^*′′*^ | ***x***_*•*_) d*k*^*′′*^ is the expected knowledge of a random adult at cultural equilibrium in the ***x***_*•*_-lineage, which we denote as 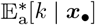. Using this notation and eq. (B.12), and factorizing by 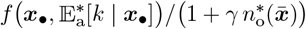, eq. (B.11) reduces to

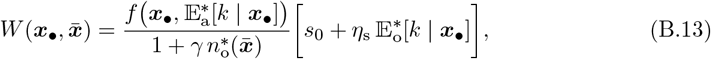

where

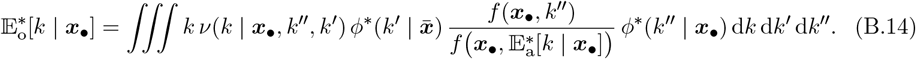

The term 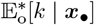 is the expected knowledge of a random offspring once it has completed learning within the ***x***_*•*_-lineage at cultural equilibrium, taking into account the probability density of parental knowledge 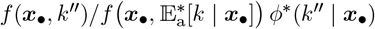, the probability density of knowledge among potential oblique exemplars 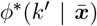, and the probability density of stochastic learning outcomes *ν*(*k* | ***x***_*•*_, *k*^*′′*^, *k*^*′*^). Note that the probability density of parental knowledge is proportional to *f* (***x***_*•*_, *k*^*′′*^)*/f* ***x***_*•*_, 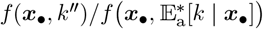 because lineage members with higher fecundity are overrepresented among parents and, consequently, among vertical exemplars.

By replacing 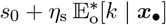 by 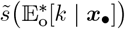 (obtained using eq. (4) with 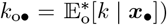 we obtain

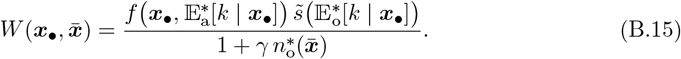

By substituting 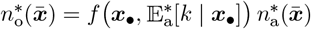 into eq. (B.15) we obtain

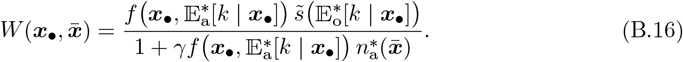

To move forward, we derive an expression for 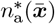. At demographic equilibrium, each adult, on average, produces one viable offspring to replace itself, so the mean fitness in the population remains equal to 1. When trait variances are small, the mean fitness can be approximated by 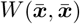, so 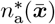 is characterized by

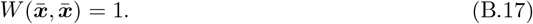

By substituting the expression for 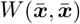 from eq. (B.16) with 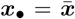 solving for 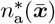 into eq. (B.17) and we obtain

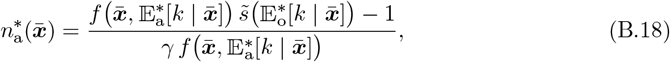

where 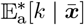 and 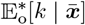 are the expected knowledge at cultural equilibrium for a random adult and a random offspring after learning is complete in an 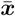-lineage. Because trait variances are small, 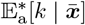 and 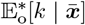 are equal to the mean adult and offspring knowledge in the population at cultural equilibrium.

By substituting the expression for 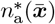 from eq. (B.18) into eq. (B.16), we obtain

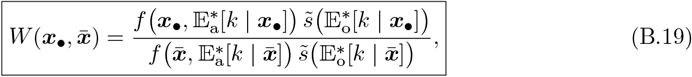

which is eq. (5) of the main text.

### B.2 Learning traits effect on lineage knowledge

Here, we derive the conditions that 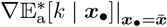 and 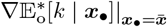 satisfy.

We first derive eq. (14) from the main text, which give the conditions that 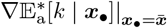 satisfies. Differentiating both sides of eq. (A.56) with respect to the traits ***x***_*•*_ and evaluating at 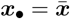, we obtain

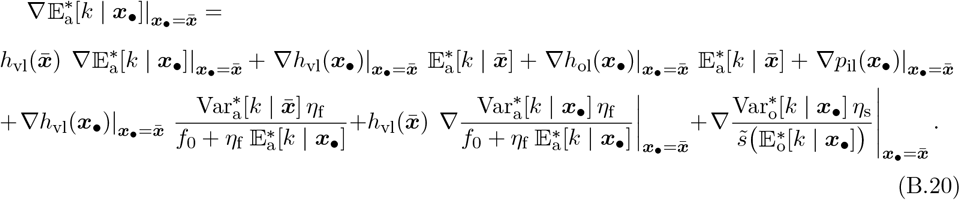

Equation (B.20) can be rewritten as

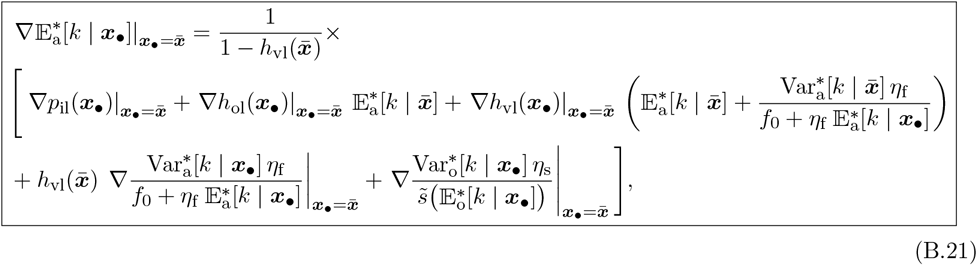

which is eq. (14) of the main text.

We now derive the conditions that 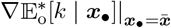 satisfies. Differentiating both sides of eq. (A.55) with respect to the traits ***x***_*•*_ and evaluating at 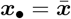, we obtain

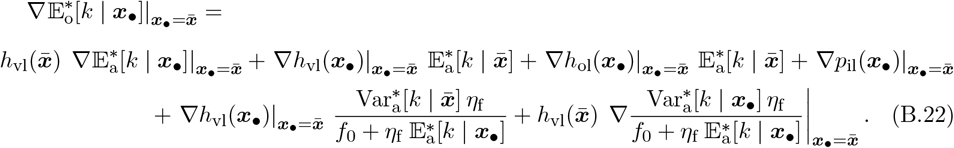

By substituting the expression of 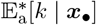 from the first line of eq. (A.54) into eq. (B.22) and rearranging we obtain

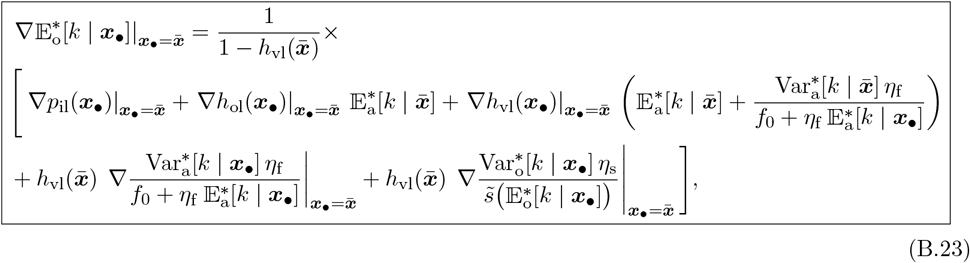

which gives a condition satisfied by 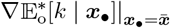.

We next link the conditions satisfied by 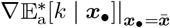 and 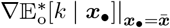. To this end, we rewrite (B.23) in terms of the expression for 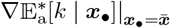 given in eq. (B.21)

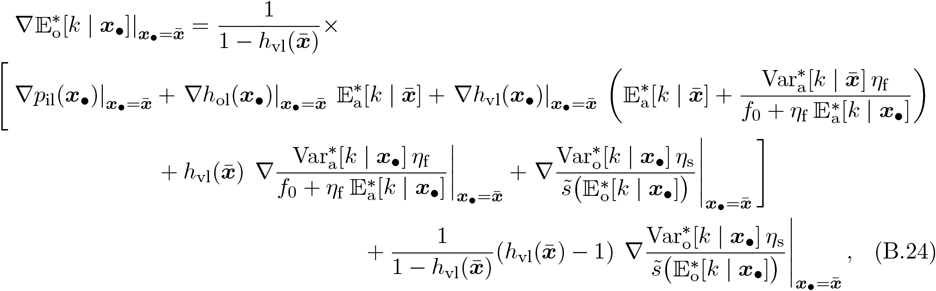

which simplifies to

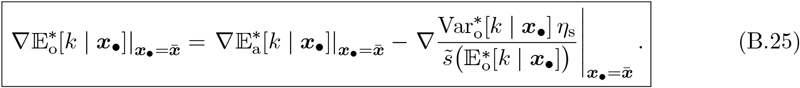

### B.3 Numerical estimation of the average learning traits favored by selection

Let 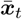 denote the mean trait values in generation *t*. The expected change in these mean trait values from one generation to the next is given by

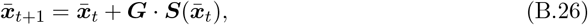

where ***G*** is the traits covariance matrix, assumed to be constant here (Iwasa et al., 1991; Mullon and Lehmann, 2019).

To numerically determine the mean trait values favored by selection 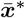, we first need to obtain the full expression for the selection gradient 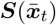. This requires deriving an exact expression 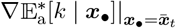 and 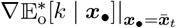.

To simplify notation, we rewrite eq. (A.54) in functional form

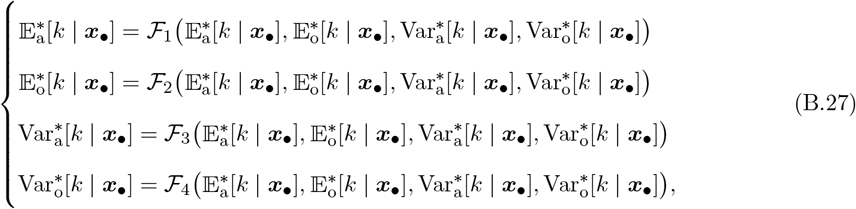

where the functions ℱ_1_, ℱ_2_, ℱ_3_ and ℱ_4_ are defined by the right-hand sides of eq. (A.54). Thus, by differentiating both sides of each line in the system (B.27) with respect to ***x***_*•*_ and evaluating at 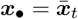, we obtain the system

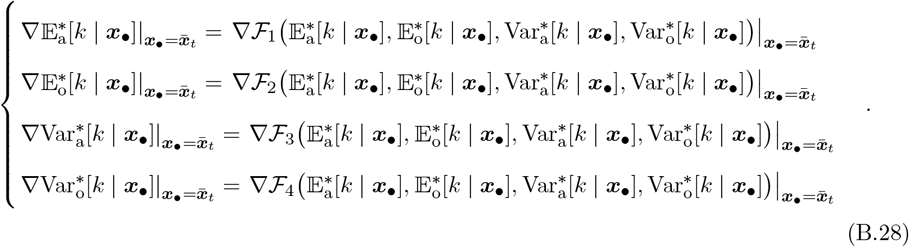

By solving eq. (B.28) for 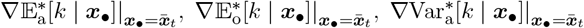 and 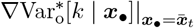 using the *Solve* function in Wolfram Mathematica 13.0.0, we obtain an exact expression for 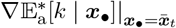 and 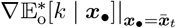 . These results enable us to derive an exact expression for the selection gradient 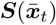 (see the Mathematica Notebook in the supplementary material).

With the expression of the selection gradient in hand, we can now numerically track the evolution of mean traits across generations. We iterated the dynamics described by eq. (B.26) with ***G*** = 0.1***I*** where ***I*** is the identity matrix, starting from the initial conditions 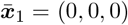 and 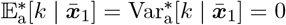. For each subsequent generation 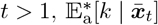 and 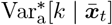 are recalculated using the procedure described in section A.4.4, setting 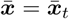 and initializing the procedure with the equilibrium values of 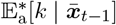 and 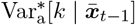 from the previous generation. We iterate this dynamics until 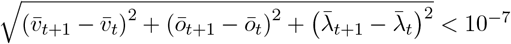.

Once the mean traits converge to an interior singular point 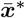 at which directional selection vanishes (i.e., 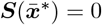), selection may become disruptive, potentially leading to evolutionary branching whereby trait distribution becomes bimodal (Geritz et al., 2016). To assess we compute the Hessian matrix

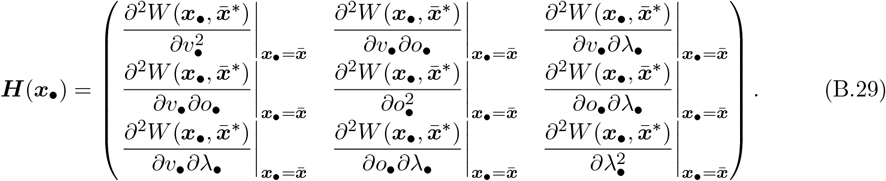

If the leading eigenvalue of ***H***(***x***_*•*_) is negative, selection is stabilizing and evolutionary branching cannot occur. In all cases considered, we find that the leading eigenvalue of ***H***(***x***_*•*_) is negative, confirming the absence of evolutionary branching.

Numerically evaluating ***H***(***x***_*•*_) requires second derivatives of 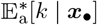 and 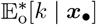 with respect to traits at 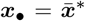. These are obtained by differentiating system (B.27) twice with respect to traits ***x***_*•*_ at 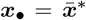 and solving the resulting system using the *Solve* function in Wolfram Mathematica 13.0.0.

### B.4 Link to the findings of Maisonneuve et al. (2025)

Here, we present the connection with the findings of Maisonneuve et al. (2025). Maisonneuve et al. (2025) showed that oblique learning can evolve even if vertical learning is more efficient, because individuals can access knowledge from other adults that differs from their parents. Oblique learning evolves when the parameter *ρ*, defined in Maisonneuve et al. (2025) as the probability that two individuals from the same generation produce simultaneously different knowledge during individual learning, is low, thereby limiting overlap in adult knowledge. In our model, we explicitly control the degree of knowledge overlap among adults using the parameter *ρ* as well. Although *ρ* is not defined identically in both models, in both cases, low values of *ρ* correspond to low overlap in the knowledge held by adults. Consistent with previous findings, we show that when adult knowledge does not fully overlap (i.e., low value of *ρ*), oblique learning can evolve, as it enables individuals to acquire information unavailable from their parents.

## Appendix C: Selection pressure on stochasticity in learning

In this section, we introduce and derive a condition for the emergence of an additional trait *ζ* ∈ [0, +∞) that affects stochasticity in individual learning *σ*_*i*_(*ζ*). We assume that 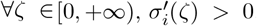, meaning that higher values of *ζ* correspond to greater stochasticity in learning. We also assume that *σ*_*i*_(0) *>* 0, with *σ*_*i*_(0) remaining small, reflecting that learning is inherently stochastic even in the absence of a specific trait promoting it.

Each individual is now characterized by a vector of traits ***x***_*•*_ = (*v*_*•*_, *o*_*•*_, *λ*_*•*_, *ζ*_*•*_). We study how selection may favor the emergence of the trait *ζ* that increases stochasticity in learning, starting from a population with mean trait vector 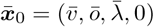. A trait located at its lower boundary will emerge if selection favors an increase in its population mean value, that is, if the corresponding component of the selection gradient is positive. Using eq. (13) we obtain that the trait *ζ* emerge if

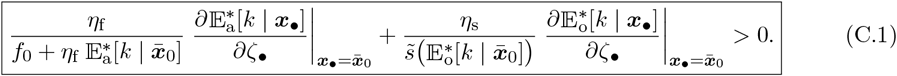

To assess the emergence of *ζ*, we need to determine in the ***x***_*•*_-lineage the effect of *ζ*_*•*_ on the expected knowledge of adults and offspring. Using eqs. (B.21) and (B.23), we find that

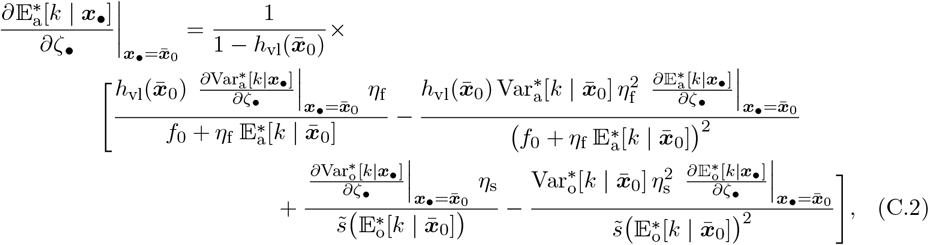

and

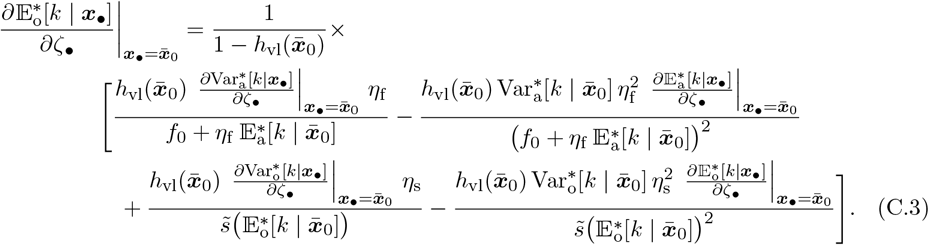

Solving eqs. (C.2) and (C.3) for 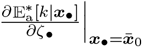 and 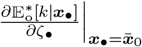 we obtain

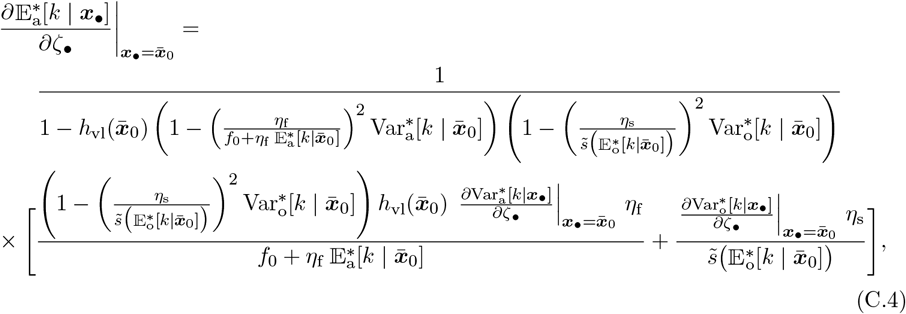

and

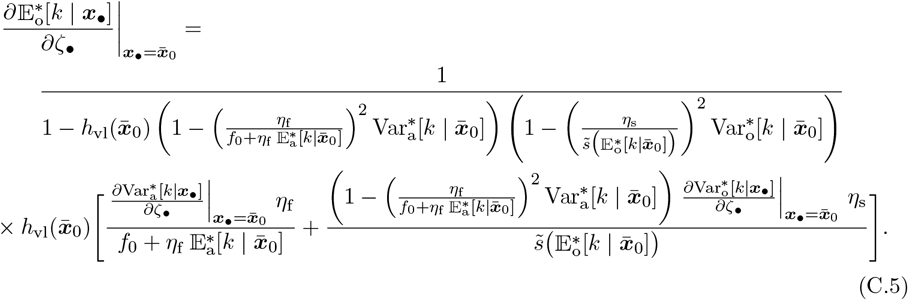

Since *σ*_i_(0) is small at 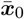, the variance in knowledge of both adults and offspring is small in each lineage. In this regime, eqs. (C.4) and (C.5) simplify to

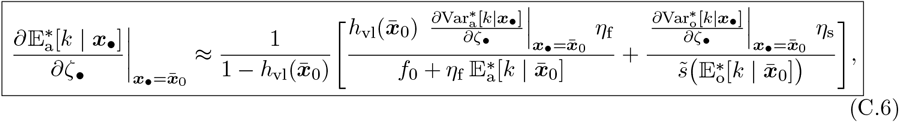

and

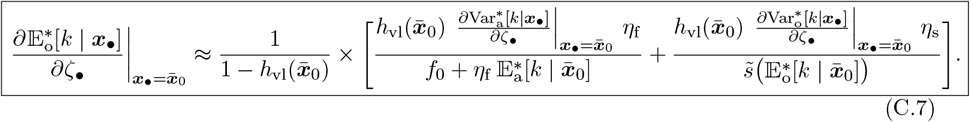

To proceed further, we determine the effect of *ζ*_*•*_ on the variance of knowledge held by a randomly chosen adult and offspring within the ***x***_*•*_-lineage. By differentiating the third and fourth lines of eq. (A.54) with respect to *ζ*_*•*_ and evaluating at 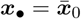 we obtain

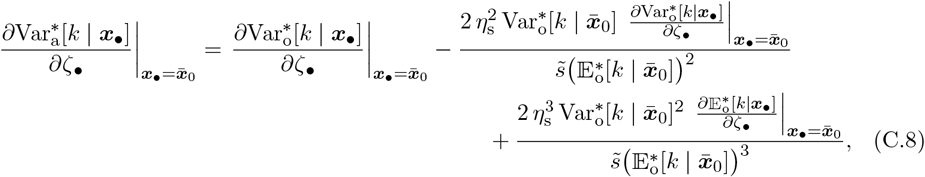

and

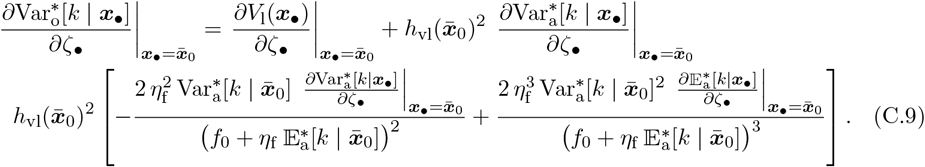

By substituting the expression of 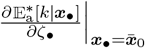 and 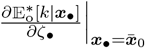 from eqs. (C.6) and (C.7) into eqs. (C.8) and (C.9) and by solving for 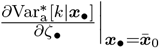 and 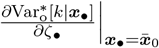 we obtain

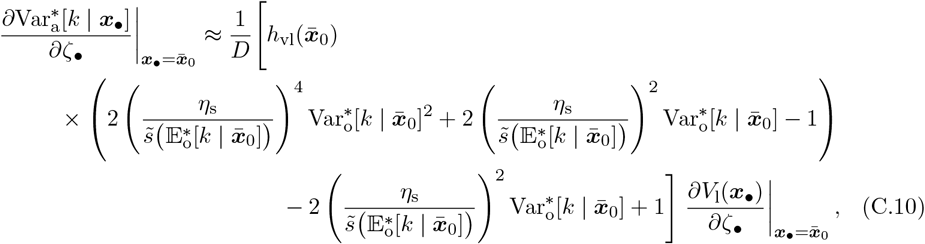

and

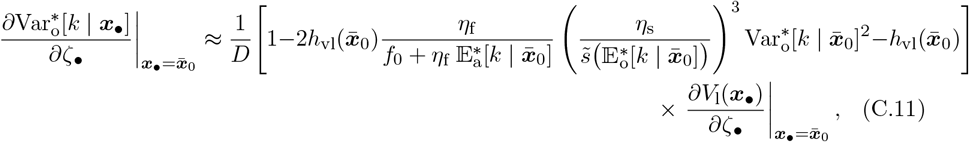

where

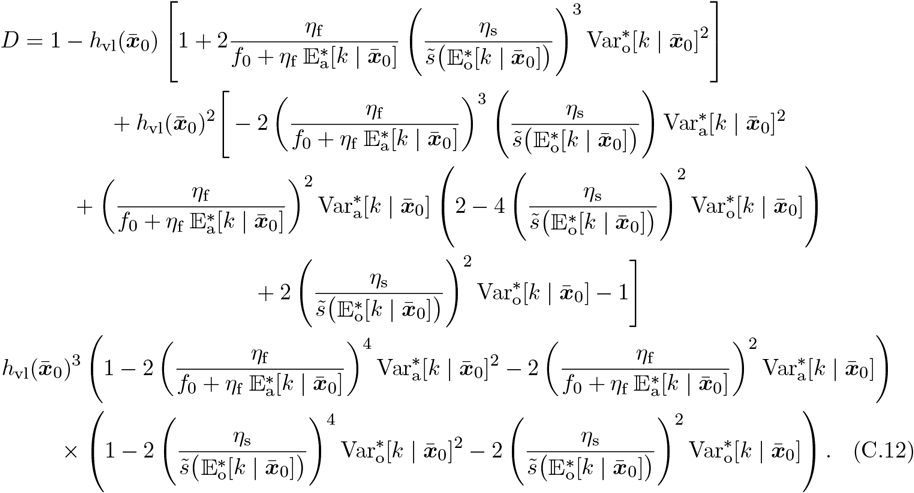

Given that the variance in knowledge of both adults and offspring is small, and using the expression of *V*_l_(***x***_*•*_) from eq. (A.47) with *σ*_i_ = *σ*_i_(*ζ*_*•*_), we obtain

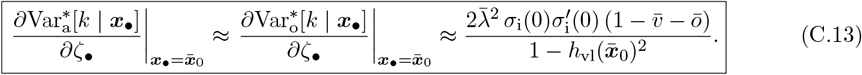

Since the right-hand side of eq. (C.13) is always positive, it shows that, in the ***x***_*•*_-lineage, the trait *ζ*_*•*_ consistently increases the variance of knowledge among randomly chosen adults and offspring within the lineage. Moreover, from eqs. (C.6) and (C.7), we infer that *ζ*_*•*_ also consistently increases their expected knowledge. Consequently, eq. (C.1) always holds, and selection systematically favors the emergence of the trait *ζ*.

## Appendix D: Individual-based simulations

Our individual-based simulations track the evolution of the trait distribution, the distribution of individual knowledge, and the population size at each generation for a fixed number of generations following the life cycle described in the main text. Each individual *i* at each generation is characterised by the vector of its three traits (*v*_*i*_, *o*_*i*_, *λ*_*i*_), as well as its knowledge *k*_*i*_ at the end of the learning period, the knowledge of its parent *k*_p,i_ and its oblique exemplar *k*_a,i_. At the first generation, the values of *v*_*i*_, *o*_*i*_, *λ*_*i*_, *k*_p,i_ and *k*_a,i_ are by default all initialized to 0 for each individual *i* ∈ {1, …, *n*_1_}, where *n*_1_ = 100 is the number of individuals generation 1.

At each generation *t* ≥ 1, the following occurs:

*(i) Reproduction*. We first determine the fecundity *f*_*i*_ of each individual *i* ∈ {1, …, *n*_*t*_}, where *n*_*t*_ is the number of adult individuals at generation *t*. The fecundity of each individual *i*, is calculated using *f*_*i*_ = *f* ((*v*_*i*_, *o*_*i*_, *λ*_*i*_), *k*_*i*_), where the expression of *f* ((*v*_*i*_, *o*_*i*_, *λ*_*i*_), *k*_*i*_) is given in eq. (1) with ***x***_*•*_ = (*v*_*i*_, *o*_*i*_, *λ*_*i*_) and *k*_p*•*_ = *k*_*i*_. Each individual *i* then produces a number of offspring that is sampled from a Poisson probability density with mean equal to *f*_*i*_. In total, they produce a total number of offspring that we denote as *n*_o,*t*_.

*(ii) Mutation*. With probability 1 − *µ*, an offspring inherits the same traits as its parent. Otherwise, with probability *µ*, traits mutate; we model this by adding an effect sampled from a normal probability density with mean 0 and variance *σ*^2^ to each parental trait value. If necessary, the resulting trait values are truncated to remain in {(*v, o, λ*) : 0 ≤ *v* ≤ 1, 0 ≤ *o* ≤ 1, 0 ≤ *v* + *o* ≤ 1, 0 ≤ *λ* ≤ 1}. In all simulations, we set *µ* = 0.01 and *σ* = 0.01.

*(iii) Learning*. For each offspring, *i*, an oblique exemplar is randomly sampled from the adults, with its knowledge denoted as *k*_a,i_. The knowledge *k*_*i*_ of each offspring *i* after learning is complete, for *i* ∈ {1, …, *n*_o,*t*_}, is determined by *k*_*i*_ = *k*_*•*_(1) where *k*_*•*_(1) is computed by solving numerically the stochastic differential equation eq. (2) with ***x***_*•*_ = (*v*_*i*_, *o*_*i*_, *λ*_*i*_), *k*_p*•*_ = *k*_p,i_ and *k*_a*•*_ = *k*_a,i_ using the *sdeint*.*itoint* function from the Python package *sdeint*. To reduce computation time when *σ*_v_ = *σ*_o_ = 0, we instead compute *k*_*•*_(1) using eq. (A.6) with ***x***_*•*_ = (*v*_*i*_, *o*_*i*_, *λ*_*i*_), *k*_p*•*_ = *k*_p,i_ and *k*_a*•*_ = *k*_a,i_, which provides an explicit expression for the final knowledge after learning, thereby avoiding numerical integration of the stochastic differential equation eq. (2).

*(iv) Survival*. All parents die and each offspring *i* in {1, …, *n*_o,*t*_} then survives till adulthood with probability *s*(*k*_*i*_, *n*_o,*t*_) (and otherwise dies). The survival probability of each offspring *i* is calculated using the expression of *s*(*k*_*i*_, *n*_o,*t*_) given by substituting the expression of 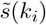 from eq. (4) with *k*_o*•*_ = *k*_*i*_ into eq. (3) with *k*_o*•*_ = *k*_*i*_ and *n*_o_ = *n*_o,*t*_. This results in the number *n*_*t*+1_ of adults of the next generation (with *n*_*t*+1_ ≤ *n*_o,*t*_).

We repeat steps (i)-(iv) for a fixed number of generations (see figure legends for parameter values).

## E Supplementary figures

**Figure S.1:**
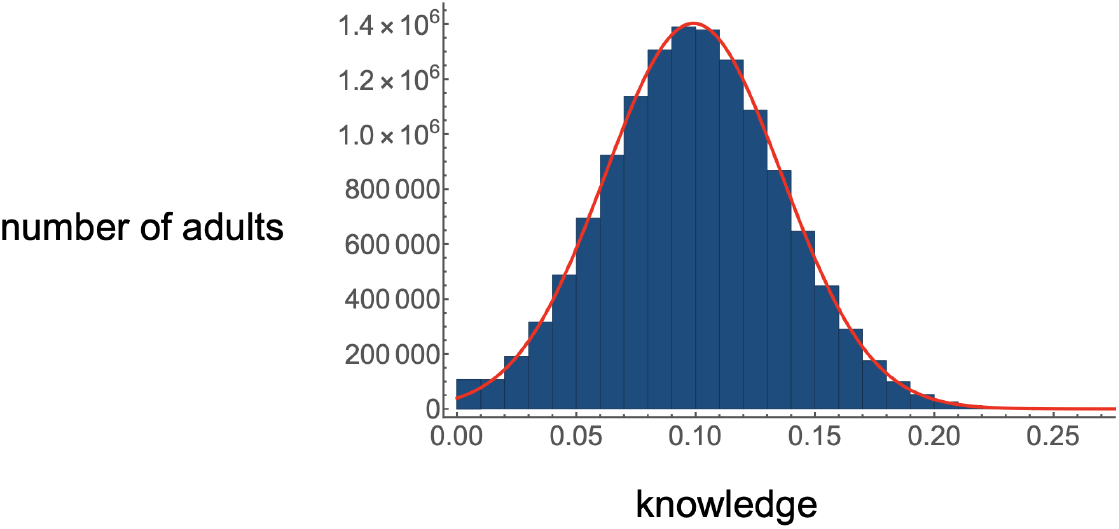
Stationary distribution of adults knowledge from individual-based simulation. The histogram shows the distribution of adult knowledge over 100,000 generations at equilibrium, based on individual-based simulations. The population was first allowed to evolve for 100,000 generations to ensure that both cultural and evolutionary equilibria were reached. The red curve represents the distribution that would be expected if adult knowledge were normally distributed, with the mean and variance of the Gaussian distribution matching those observed in the simulation, illustrating that a Gaussian approximation provides a good fit for the distribution of knowledge in the population. Parameters are: *f*_0_ = 5, *s*_0_ = 1, *β*_v_ = 1.4, *β*_o_ = 1.3, *α* = 0.1, *ϵ* = 0.05, *ρ* = 0.05, *σ*_v_ = *σ*_o_ = 0, *σ*_i_ = 0.05, *η*_f_ = 25, *η*_s_ = 5, *γ* = 0.01, *θ* = 0.1.

**Figure S.2:**
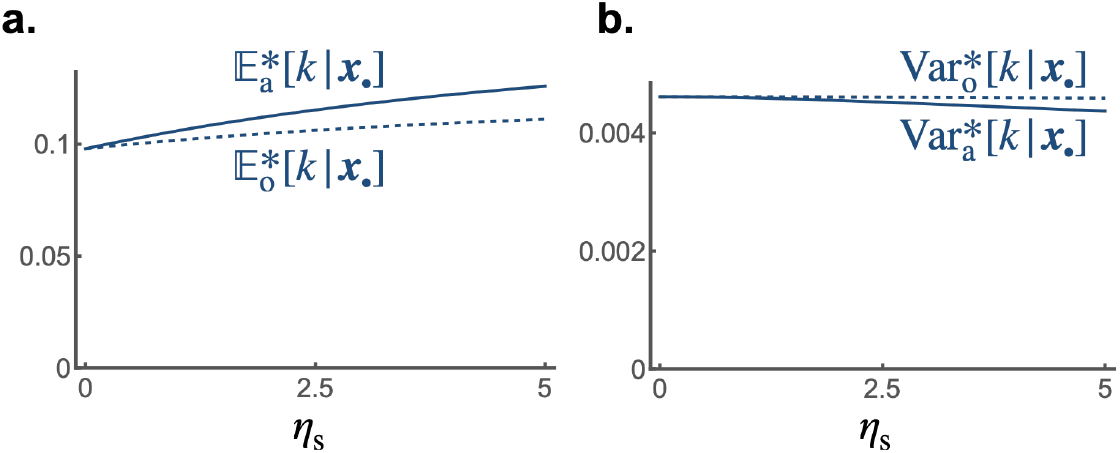
The impact of the conversion factors that translate knowledge into survival benefits *η*_s_ on the expected knowledge and variance in knowledge at cultural equilibrium for a random adult and offspring of the *x*_*•*_-lineage. **a** 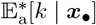 (blue solid line) and 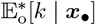 (blue dashed line) and **b** 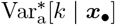 (blue solid line) and 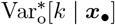 (blue dashed line) according to *η*_s_. This shows that the expected knowledge in adults is slightly higher than in offspring, while the variance in adult knowledge is slightly lower than in offspring. Default parameters are the same as in fig. 2 with 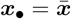.

**Figure S.3:**
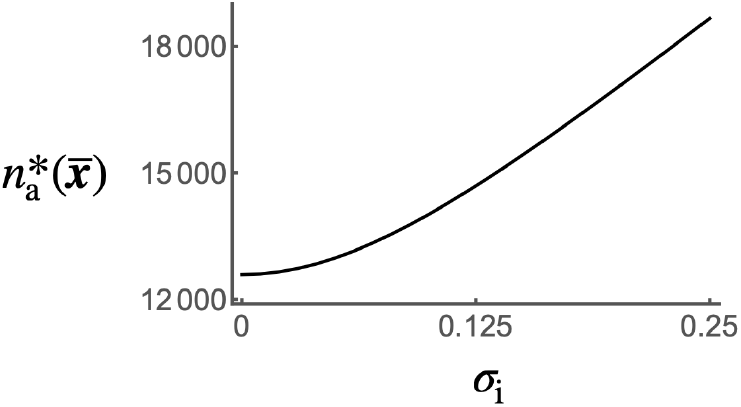
The impact of stochasticity in individual learning on population size. Equilibrium adult population size 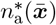 according to *σ*_i_. This shows that the population size increases with stochasticity in individual learning. Default parameters are the same as in fig. 2 with *γ* = 10^−4^.

**Figure S.4:**
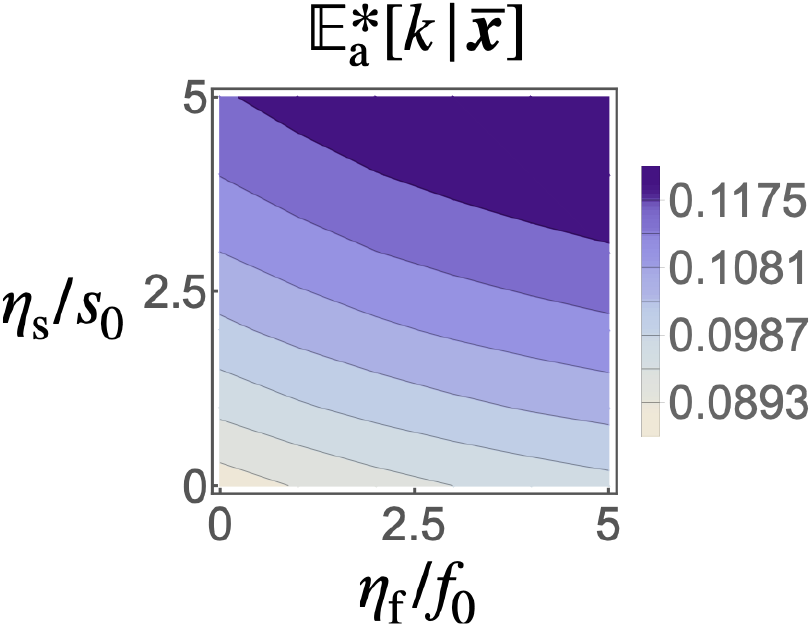
Impact of selective pressure on the population mean knowledge. Population mean knowledge at cultural equilibrium 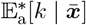 according to *η*_f_ and *η*_s_. This shows that the effect of selection in enhancing the mean knowledge 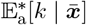 is stronger when *η*_f_ and *η*_s_ are higher. Default parameters are the same as in fig. 2.

**Figure S.5:**
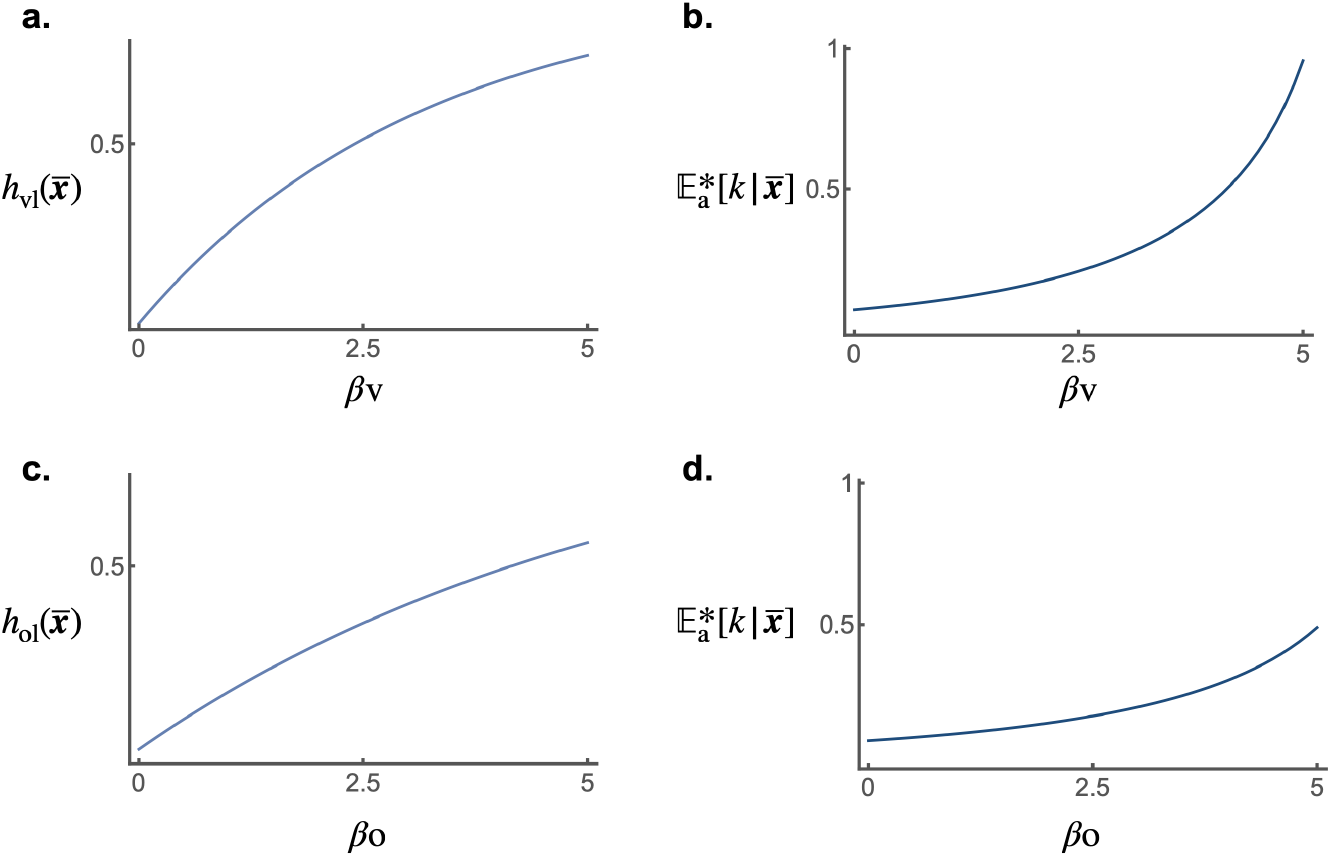
Impact of transmission on the population mean knowledge. Population mean **a** vertical cultural heritability 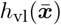 and **b** knowledge at cultural equilibrium 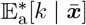 according to *β*_v_. Population mean **c** oblique cultural heritability 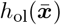 and **d** knowledge at cultural equilibrium 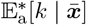 according to *β*_o_. This shows that the effect of selection in enhancing the mean knowledge 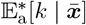 is stronger when intergenerational transmission is higher. Default parameters are the same as in fig. 2.

**Figure S.6:**
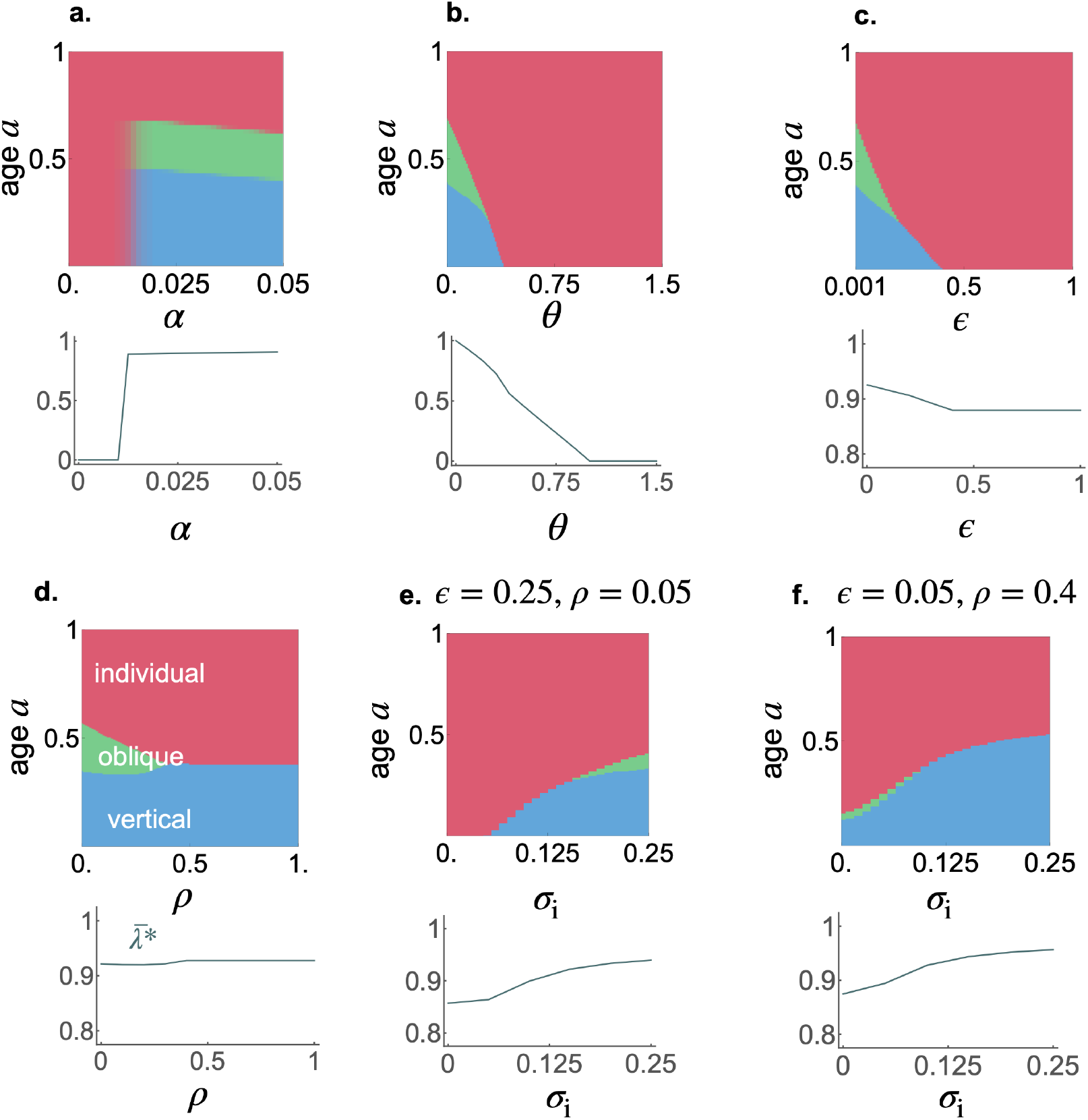
Factors impacting the evolution of learning traits. Learning schedule and investment in learning 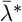 at 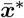 against **a** *α*, **b** *θ*, **c** *ϵ*, **d** *ρ*, and **e-f** *σ*_i_. Blue, green, and pink areas represent time spent performing vertical, oblique, and individual learning, respectively. Default parameters are the same as in fig. 3, except in panel e, where *ϵ* = 0.25 and in panel f, where *ρ* = 0.4.

**Figure S.7:**
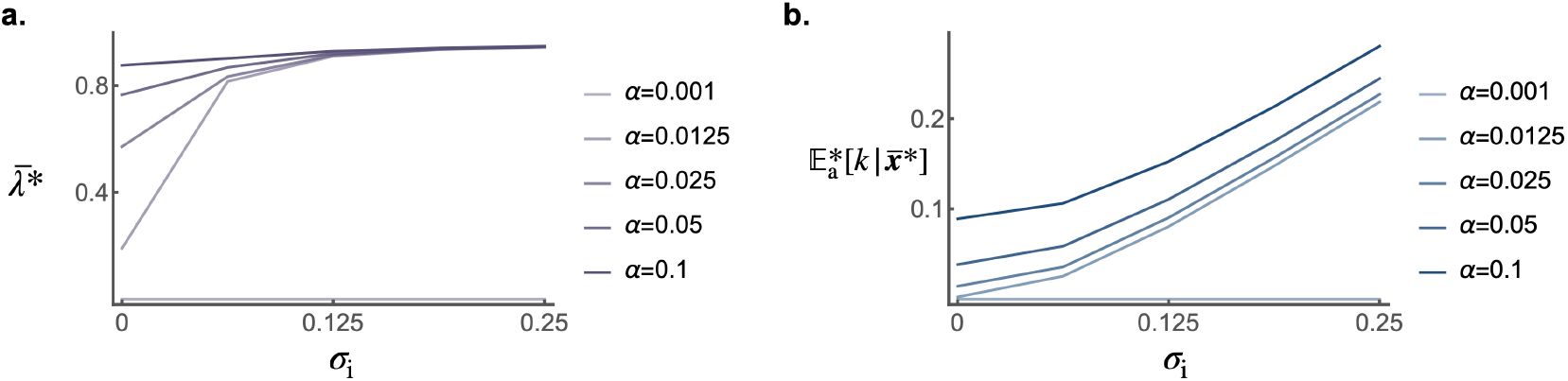
Impact of stochasticity in individual learning *σ*_i_ on investment in learning 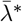 and the population mean knowledge 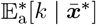 at 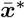 for different values of *α*. This shows that stochasticity in learning can promote both higher average knowledge and greater investment in learning, except in populations with very low values of *α*. Default parameters are the same as in fig. 3.

**Figure S.8:**
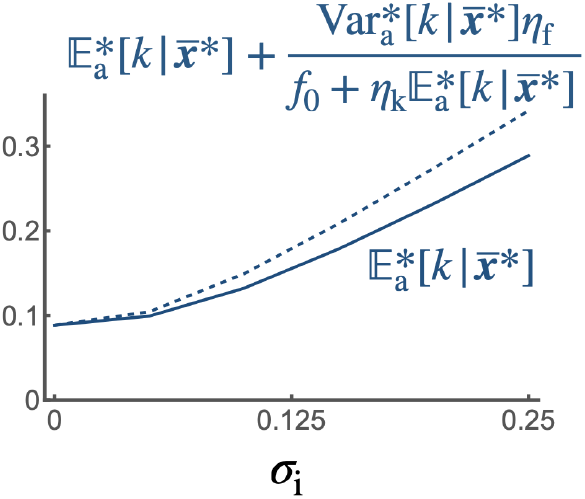
Impact of stochasticity in individual learning *σ*_i_ on the average knowledge of parents 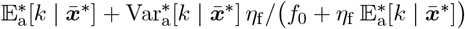 and of oblique exemplars 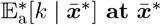. This shows that under higher stochasticity in individual learning, the average knowledge of parents increases more than that of oblique exemplars. Default parameters are the same as in fig. 3.

**Figure S.9:**
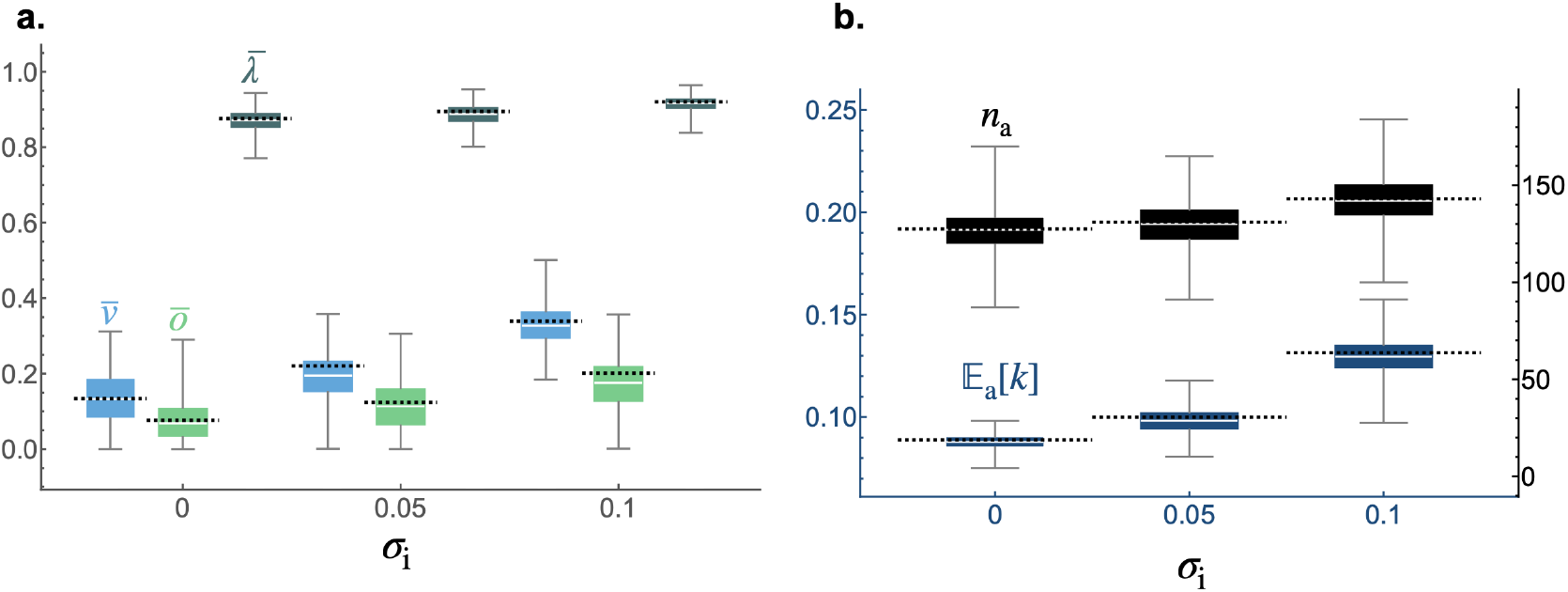
Impact of stochasticity in individual learning on the stationary distribution of mean traits, average adults knowledge, and adult population size in individual-based simulations. Box plots show distributions of **a** mean traits 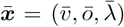 and **b** average adults knowledge 𝔼_a_(*k*) and adult population size *n*_a_ over 400,000 generations at equilibrium from individual-based simulations for different values of the strength *σ*_i_ of stochasticity in individual learning. For each value of *σ*_i_, the population was first allowed to evolve for 100,000 generations to ensure equilibrium was reached. The box shows the interquartile range (IQR) with the median marked inside; whiskers extend to the smallest and largest values within 1.5 × IQR from Q1 and Q3, respectively. Dashed lines show the predicted values derived from evolutionary analyses. The close match between simulation outcomes and analytical predictions suggests a good level of agreement between the two approaches. Default parameters are the same as in fig. 3 with *σ*_v_ = *σ*_o_ = 0 and *γ* = 0.01.

**Figure S.10:**
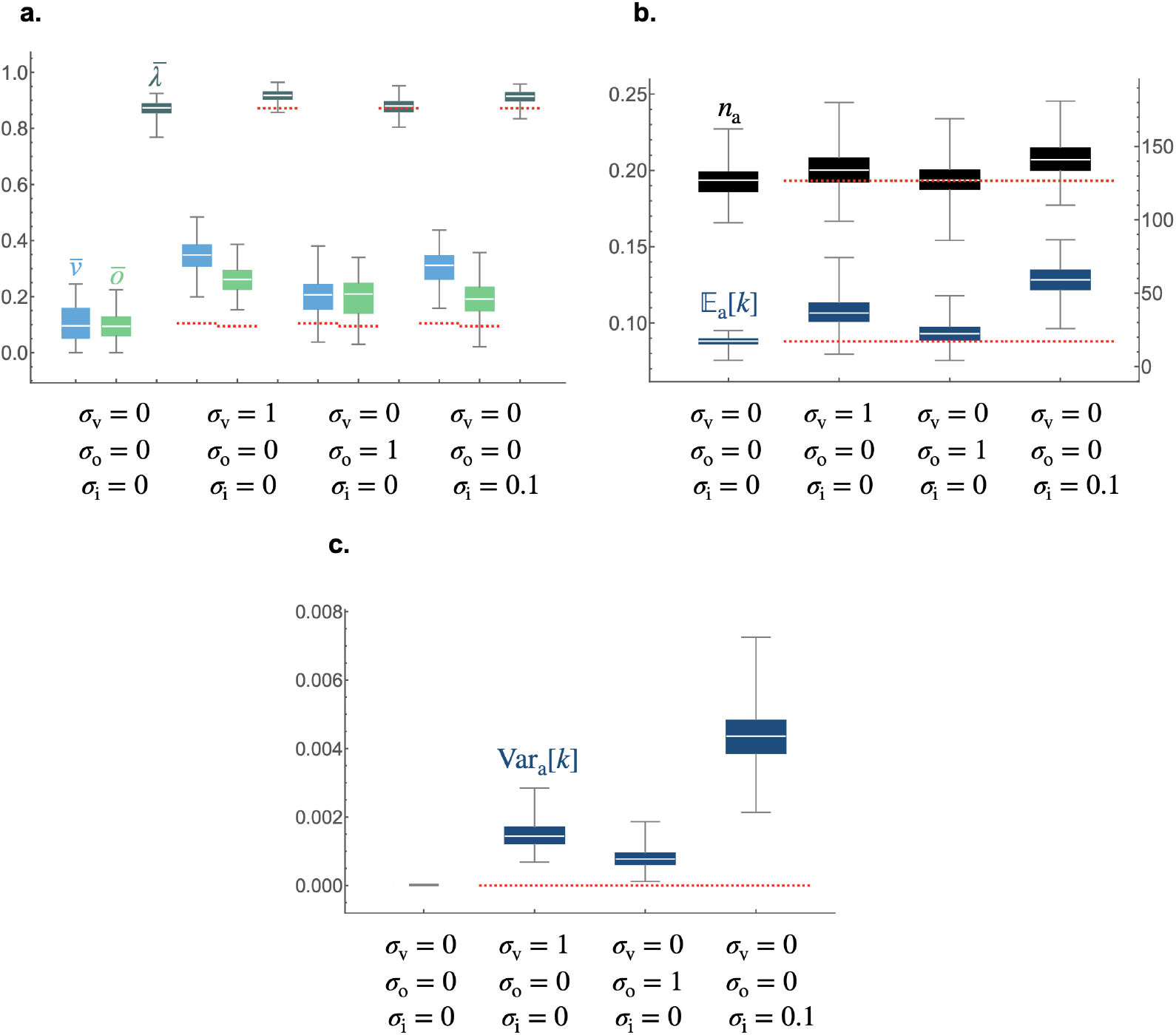
Impact of stochasticity in the different types of learning on the stationary distribution of mean traits, average adults knowledge, and adult population size in individual-based simulations. Box plots show distributions of **a** mean traits 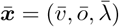, **b** average adults knowledge 𝔼_a_(*k*) and adult population size *n*_a_ and **c** adults knowledge variance Var_a_(*k*) over 100,000 generations at equilibrium from individual-based simulations for different values of strength of stochasticity in vertical *σ*_v_, oblique *σ*_o_, and individual *σ*_i_ learning. For each combination of values for *σ*_v_, *σ*_o_, and *σ*_i_, the population was first allowed to evolve for 100,000 generations to ensure equilibrium was reached. Because individual-based simulations run significantly slower when *σ*_v_ *>* 0 or *σ*_o_ *>* 0 (due to the need to numerically solve the stochastic differential equation eq. (2)), we initialize trait values for each individual *i* in the first generation as (*v*_*i*_, *o*_*i*_, *λ*_*i*_) = (0.13404, 0.0762715, 0.876229). These correspond to the mean trait values predicted at 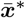 in the absence of stochasticity in learning (i.e., when *σ*_v_ = *σ*_o_ = *σ*_i_ = 0). The box shows the interquartile range (IQR) with the median marked inside; whiskers extend to the smallest and largest values within 1.5 × IQR from Q1 and Q3, respectively. Red dashed lines show the mean values obtained for individual-based simulation in the absence of stochasticity in learning. This shows that, similarly to individual learning, greater stochasticity in vertical and oblique learning promotes the accumulation of knowledge, an increase in population size, and a greater allocation of time to social learning. Note that these effects are less pronounced for stochasticity in oblique learning, because it generates less variance in knowledge and therefore weaker cultural selection. This likely occurs because the amount of knowledge acquired through oblique learning is lower, so stochasticity produces less variation overall. Default parameters are the same as in fig. 3 with *γ* = 0.01.

## Notes

https://github.com/Ludovic-Maisonneuve/stochasticity_evolution_learning

### Competing Interest Statement

The authors have declared no competing interest.

### Summary of Updates

We corrected typos in the mansucript

